# GAK and PRKCD are positive regulators of PRKN-independent mitophagy

**DOI:** 10.1101/2020.11.05.369496

**Authors:** Michael J. Munson, Benan J. Mathai, Laura Trachsel, Matthew Yoke Wui Ng, Laura Rodriguez de la Ballina, Sebastian W. Schultz, Yahyah Aman, Alf H. Lystad, Sakshi Singh, Sachin Singh, Jørgen Wesche, Evandro F. Fang, Anne Simonsen

## Abstract

The mechanisms involved in programmed or damage-induced removal of mitochondria by mitophagy in response to different stimuli remains elusive. Here, we have screened for regulators of PRKN-independent mitophagy using an siRNA library targeting 197 proteins containing lipid interacting domains. We identify Cyclin G-associated kinase (GAK) and Protein Kinase C Delta (PRKCD) as novel regulators of PRKN-independent mitophagy, with both being dispensable for PRKN-dependent mitophagy and starvation-induced autophagy. We demonstrate that the kinase activity of both GAK and PRKCD are required for efficient mitophagy *in vitro*, that PRKCD is present on mitochondria, and that PRKCD is required for ULK1/ATG13 recruitment to early autophagic structures. Importantly, we demonstrate *in vivo* relevance for both kinases in the regulation of basal mitophagy. Knockdown of GAK homologue (gakh-1) in *C.elegans* or PRKCD homologues in zebrafish led to significant inhibition of basal mitophagy, highlighting the evolutionary relevance of these kinases in mitophagy.

## INTRODUCTION

The selective degradation of mitochondria by autophagy (mitophagy) is important for cellular homeostasis and disease prevention. Defective clearance of damaged mitochondria is linked to the development of neurodegenerative diseases such as Parkinson’s disease (PD) and has also been linked to cancer^1^. Damaged mitochondria have the potential to leak dangerous reactive oxygen species causing cell damage and ultimately death, so their rapid clearance is favoured. Mitophagy involves the sequestration of mitochondria into double-membrane structures termed autophagosomes that transport and deliver material to the lysosome for degradation^2^. Elucidation of the molecular mechanisms of mitophagy has largely focused upon hereditary forms of PD and the role of the genetic risk genes *PINK1* and *Parkin* (*PRKN*) in mediating mitophagy in response to mitochondrial depolarisation ^3^. Such studies have shown that selective recognition of damaged mitochondria involves PINK1-mediated phosphorylation of ubiquitin and PRKN. This further ubiquitinates outer mitochondrial membrane proteins that are recognised by specific ubiquitin-binding autophagy receptors that interact with LC3 and GABARAP proteins in the autophagy membrane to facilitate mitophagosome formation. The relative importance of this pathway *in vivo* is however not clear, as the vast majority of basal mitophagy occurring *in vivo* seems largely independent of PINK1/PRKN, as demonstrated in both mice and fly models ^4,5^. Consequently, further characterisation of the mechanisms involved in PRKN-independent basal mitophagy pathways is needed to understand their role in normal physiology and disease development. One such pathway is the HIF1α/hypoxia-dependent pathway that has been particularly well characterised for the clearance of red blood cell mitochondria that occur via upregulation of the mitophagy receptor BNIP3 ^6^. Several small molecules have been identified to stabilise HIF1α and replicate a hypoxia-induced mitophagy phenotype without the requirement for hypoxic conditions ^7^, including cobalt chloride, dimethyloxaloylglycine (DMOG) and iron chelators, with deferiprone (DFP) found to be one of the most potent ^8,9^.

Whilst much progress has been made in our understanding of selective autophagy and the proteins involved, little is known about the lipids and lipid-binding proteins involved in cargo recognition and autophagosome biogenesis during selective autophagy. Formation of autophagosomes relies upon a multitude of trafficking processes to manipulate and deliver lipids to the growing structure and several proteins containing lipid interaction domains have been found to play important roles in modulating autophagy ^10^. Here we have carried out an imaging-based screen to examine whether human proteins containing lipid-binding domains have novel roles in HIF1α dependent mitophagy. We identify a shortlist of eleven novel and previously unknown candidates that regulate mitophagy. In particular, we show that the two kinases GAK and PRKCD are specifically required for HIF1α-dependent mitophagy without affecting PRKN-dependent mitophagy and that this regulation is also observed *in vivo* upon basal mitophagy. Therefore, these kinases represent novel targets for the study and regulation of basal mitophagy.

## RESULTS

### Induction and verification of mitophagy in U2OS cells

To monitor mitophagy in cultured cells, U2OS cells were stably transfected to express a tandem tag mitophagy reporter containing EGFP-mCherry fused to the mitochondrial localisation sequence (MLS) of the mitochondrial matrix protein NIPSNAP1 (hereafter termed inner MLS [IMLS] cells) in a doxycycline-inducible manner^11^. U2OS cells contain low endogenous PRKN levels that are insufficient to induce mitophagy in response to mitochondrial membrane depolarisation^12^. However, while a yellow mitochondrial network is seen under normal conditions, induction of mitophagy with the iron chelator DFP for 24 h causes movement of mitochondria to lysosomes, where the EGFP signal is quenched due to the acidic pH, leading to the appearance of red only punctate structures (Fig. 1a,b). Co-staining for the inner mitochondrial protein TIM23 verified that the EGFP-mCherry tag was localised to the mitochondrial network (Fig. 1a). To verify that the red only structures represent autolysosomes/mitolysosomes, the V-ATPase inhibitor Bafilomycin A1 (BafA1) was added for the final 2 h of DFP treatment to raise lysosomal pH and restore the EGFP signal^13^. As predicted, the number of red only structures dropped drastically (Fig. 1b). Similarly, siRNA-mediated depletion of the key autophagy inducer ULK1 before DFP addition significantly reduced the formation of red only structures (Fig. 1b). The localisation of the IMLS reporter was further validated by correlative light and electron microscopy (CLEM) following DFP treatment. Indeed, yellow network structures observed by confocal fluorescence microscopy corresponded to mitochondrial structures by EM, whereas red only structures demonstrated typical autolysosome morphologies (Fig. 1c).

**Figure 1.**
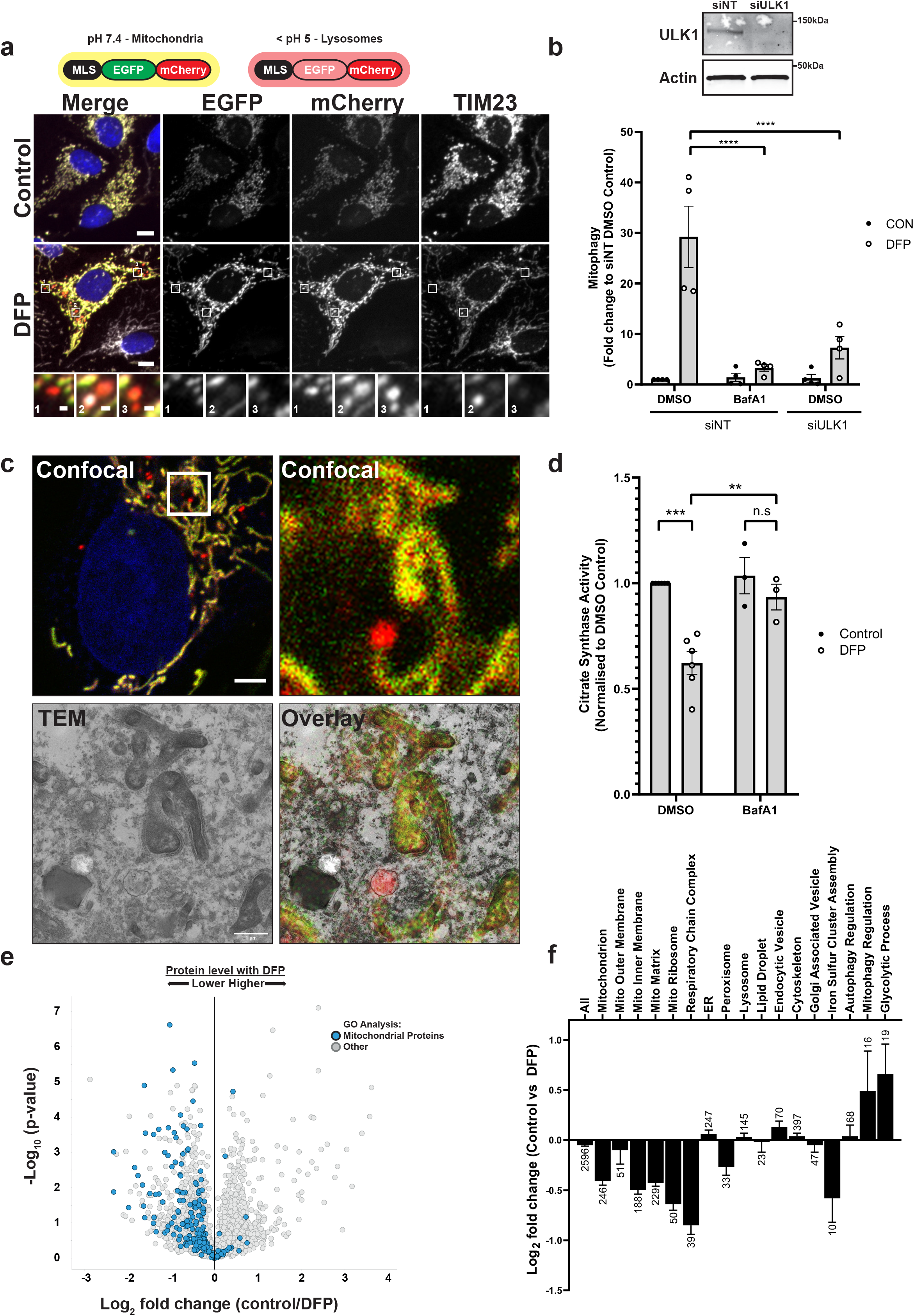
Measuring DFP-induced mitophagy in vitro. **a** U2OS cells stably expressing an internal MLS-EGFP-mCherry (IMLS) reporter that is pH responsive (yellow at neutral, red at acidic pH) were incubated for 24 h ± 1mM DFP followed by PFA fixation, antibody staining for TIM23 (Alexa Fluor-647) and widefield microscopy. Scale bar = 10 μm, inset = 0.5 μm. **b** U2OS IMLS cells were transfected with 7.5 nM siRNA non-targeting control (siNT) or siULK1 for 48 h prior to 24 h treatment ± 1 mM DFP in the presence or absence of 50 nM BafA1 for the final 2 h. Western blot from cell lysates shows representative ULK1 knockdown level. Graph represents the mean red only area per cell from fluorescence images normalised to control DMSO siNT cells ± SEM from n=4 independent experiments. Significance was determined by two-way ANOVA followed by Tukey’s multiple comparison test. **c** U2OS IMLS cells treated with 1 mM DFP as in **a** and fixed for CLEM analysis. Inset of cell area in white box is shown by confocal analysis and EM section along with EM overlay. Scale bar = 5μm, inset = 1μm **d** Citrate synthase activity from U2OS cells treated for 24 h ± 1 mM DFP with final 16 h in the presence of 50 nM BafA1 or DMSO, values are normalised to DMSO control from n=6 (DMSO) or n=3 (+BafA1) independent experiments ± SEM. Significance was determined by two-way ANOVA followed by Sidak’s multiple comparisons test. **e** U2OS whole cell protein abundance was determined by mass spectrometry following treatment ± 1 mM DFP 24 h. Mitochondrial proteins identified by GO analysis (Term = Mitochondrion) are highlighted in blue. f Mean T-test difference between Control and DFP samples for peptides identified in **e** matching GO terms related to cellular organelles. Bars represent Log2 fold change (Control vs DFP) ± SEM, number on bars indicate how many protein targets are included in GO analysis. ** = p < 0.01, *** = p < 0.001, **** = p < 0.0001 and n.s = not significant in all relevant panels.

DFP-induced mitophagy could be demonstrated biochemically by measuring the enzymatic activity of the mitochondrial matrix protein citrate synthase ^14^. Treatment with DFP for 24 h reduced citrate synthase activity by ~40 %, which could be prevented by addition of BafA1 for the final 16 h, confirming that the reduction was due to lysosomal-mediated degradation (Fig. 1d). Finally, we were able to demonstrate by proteomic analysis that the addition of DFP for 24 h decreased the abundance of multiple mitochondrial proteins (classified by gene ontology [GO] analysis) compared to control-treated cells (Fig. 1e). Comparison of different cellular organelles and compartments by GO annotation highlighted that mitochondrial proteins resident to the inner membrane and matrix were particularly reduced in response to DFP treatment (Fig. 1f, Supplementary Table 1). Peroxisomal protein abundance was also slightly reduced, whilst proteins belonging to the ER, lysosomes or endosomes were generally unaffected (Fig. 1f). In contrast, proteins involved in processes defined as glycolytic or mitophagy regulation showed increased abundance.

Taken together, we show that DFP treatment induces a lysosomal-dependent loss of mitochondrial proteins in U2OS cells and that mitolysosomes could be quantified by image analysis in U2OS IMLS cells.

### Screening for lipid-binding protein regulators of mitophagy

To uncover the mechanisms involved in selective recognition and turnover of mitochondria in DFP treated cells we carried out an image-based siRNA screen monitoring the formation of red-only structures in response to DFP following siRNA mediated knockdown of 197 putative lipid binding proteins in U2OS IMLS cells. An initial list of proteins containing established lipid interacting protein domains (FYVE, PX, PH, GRAM, C1, C2, PROPPIN, ENTH) were identified using ExPASy Prosite (see Methods and Appendix 1). This preliminary target list was cross-examined with several U2OS proteomic datasets and restricted to proteins validated to be expressed in U2OS cells ^15,16^. We included all FYVE or PX domain containing proteins due to the relevance of phosphatidylinositol 3-phosphate (PtdIns(3)P) binding proteins in autophagy initiation ^17^.

The primary screen was carried out using a pool containing three different siRNA oligonucleotides sequences per target. In the absence of DFP treatment, no spontaneous induction of mitophagy was observed upon gene knockdown (data not shown). In contrast, significant changes were observed with siRNA treatment in the presence of DFP, which are plotted as fold change relative to non-targeting (siNT) samples and grouped based upon lipid binding domains (Fig. 2a-e). As previously shown, BafA1 treatment or depletion of ULK1 strongly inhibited DFP-induced formation of red (mitolysosome) structures (Fig. 1b, Fig. 2a-e red bars). Interestingly, significantly increased levels of mitophagy were seen following knockdown of HS1BP3 (Fig. 2a), a negative regulator of starvation induced autophagy that has not previously been examined in mitophagy ^18^. It is interesting to note that relatively few hits from proteins containing a FYVE or PX domain were found compared to C1, C2 or GRAM domain containing proteins (Fig. 2f).

**Figure 2.**
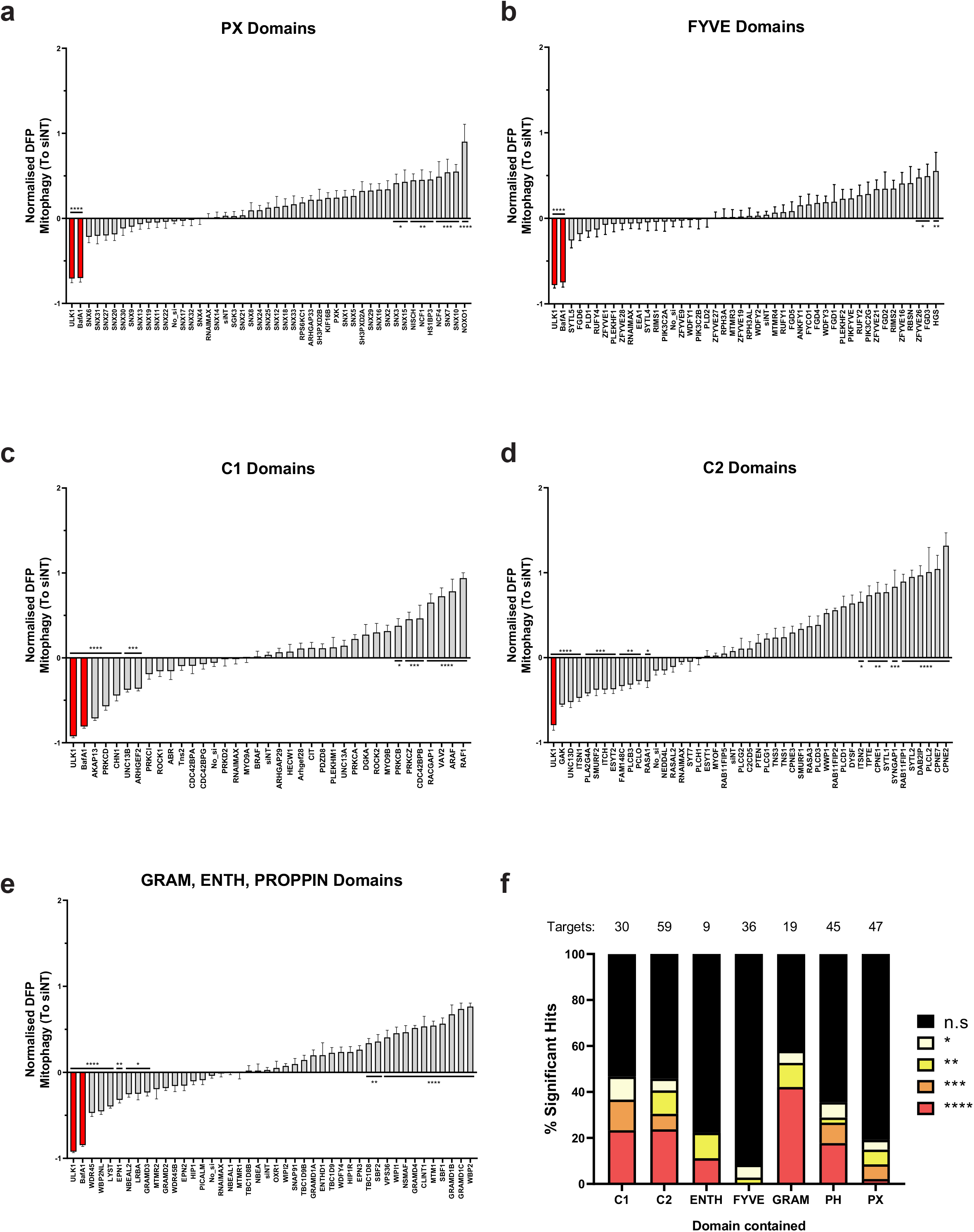
siRNA screen for lipid binding proteins involved in DFP-induced mitophagy. **a-e** Primary siRNA screen data. U2OS cells were transfected with a pool of 3 sequence variable siRNA oligonucleotides per gene target (2.5 nM per oligo) for 48 h before addition of 1 mM DFP for 24 h. Cells were PFA fixed and imaged using a 20x objective (35 fields of view per well). The red area per cell was normalised to the average of siNT controls and adjusted so that the DFP siNT control was 0 from n= 14 plates (**a**,**b**) or n=6 plates (**c-e**) ± SEM. siULK1 and BafA1 (red bars) are negative controls. siRNA targets containing similar lipid binding domains were assayed and plotted together as shown for **a** PX domains, **b** FYVE domains **c** C1 domains **d** C2 domains **e** GRAM, ENTH, PROPPIN domains. **f** Summary of significance of different lipid binding domains relative to the total tested from a e, proteins containing more than one type of domain are represented in each category. Significance was determined by one-way ANOVA followed by Dunnett’s multiple comparison test to the siNT control where * = p < 0.05, ** = p < 0.01, *** = p < 0.001, **** = p < 0.0001 and n.s = not significant in all relevant panels.

### GAK and PRKCD identified as DFP mitophagy regulators by siRNA screening

To validate prospective positive and negative regulators that demonstrated significant changes in the primary screen, we selected 29 candidates for a secondary deconvolution screen where the three siRNA oligonucleotides used in the primary screen were examined individually in U2OS IMLS cells. Their effect on DFP-induced mitophagy was analysed and quantified by high-content imaging and the level of residual target mRNA was quantified by qPCR. The mitophagy phenotype of the individual siRNA oligos was then correlated to their knock-down efficiencies and compared to the mitophagy effect of the siRNA pool used in the primary screen (Supplementary Fig. 1a). As the knockdown efficiencies for some of the targets were disappointing in the secondary screen, we carried out a tertiary screen for all targets that were found to significantly affect mitophagy in the primary screen, using individual siRNA oligos at a higher concentration of siRNA (Supplementary Fig. 1b). Based on the results of the secondary and tertiary screen, we highlighted eleven targets where at least two of the three oligos had a significant effect on DFP-induced mitophagy (Fig. 3a).

**Figure 3.**
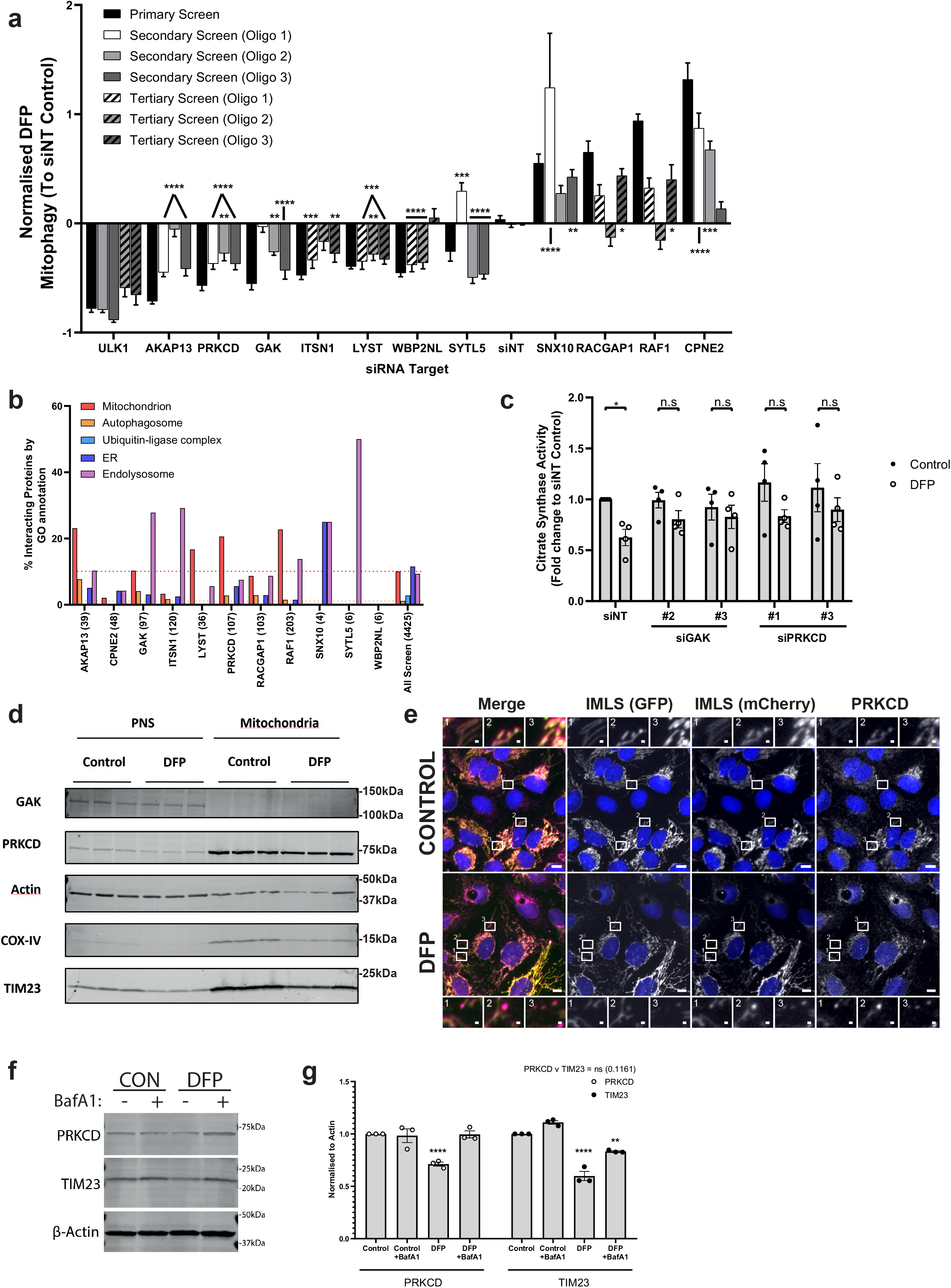
GAK and PRKCD are regulators of DFP mediated mitophagy. **a** Summary of significant targets identified across primary, secondary (7.5 nM individual siRNA oligos) and tertiary screen (15 nM each oligo) siRNA screens. Cells were transfected for 48 h prior to 24 h of 1 mM DFP treatment. Bars represent mean fold change in mitophagy relative to the siNT controls ± SEM. Significance was determined by one-way ANOVA followed by Dunnett’s multiple comparison test to the siNT control. **b** Protein-protein interaction networks for candidate proteins (see methods) were plotted by % of interacting proteins belonging to each highlighted compartment, value in brackets represents total number of interacting proteins. **c** siRNA treatment with indicated oligos for 48 h prior to 24 h treatment ± 1 mM DFP and subsequent analysis of citrate synthase activity levels. Values were normalised to the siNT control and plotted ± SEM for n=4 independent experiments. Significance was determined by two-way ANOVA followed by Sidak’s multiple comparison test. **d** U2OS cells treated ± 1 mM DFP for 24 h were enriched from a post-nuclear supernatant (PNS) for mitochondria followed by western blotting for the indicated proteins. **e** U2OS IMLS cells treated ± 1 mM DFP for 24 h followed by PFA fixation and staining for endogenous PRKCD (Alexa Fluor-647). Images were obtained by 20x objective using a Zeiss AxioObserver. Scale bar = 10 μm, insets = 1 μm. **f** U2OS cells treated ± 1 mM DFP for 24 h ± 50 nM BafA1 for the final 16 h and blotted for the indicated proteins. **g** Quantitation of PRKCD and TIM23 levels to β-actin in **f** from n=3 independent experiments ± SEM. Significance was determined by two-way ANOVA followed by Dunnett’s multiple comparison test to the control. * = p < 0.05, ** = p < 0.01, *** = p < 0.001, **** = p < 0.0001 and n.s = not significant in all relevant panels.

To identify candidates with the highest possible relevance for mitophagy, we searched for interacting proteins for each of these eleven candidate proteins using network analysis of protein interaction data (see Methods). Interacting proteins were subjected to GO analysis and grouped based upon compartments of interest, including mitochondria, autophagy and endolysosomal compartments. The percentage of interacting proteins in each group was compared to the average percentage for all proteins screened. Of interest, AKAP13 had a high number of interactors linked to mitochondria and autophagy. Additionally, GAK and PRKCD had higher numbers of autophagy and mitochondria linked proteins respectively than all screened proteins (Fig. 3b and Supplementary Fig. 2a,b). As several mitophagy receptor proteins involved in HIF1α-induced mitophagy are regulated by phosphorylation (including BNIP3 and BNIP3L), whilst the PINK1 kinase is dispensable for PRKN-independent mitophagy, we decided to further characterise the role of GAK and PRKCD kinases in mitophagy. In addition, GAK has been linked as a risk factor for PD ^19^ and PRKCD has been noted to translocate to mitochondria previously ^20^.

All three oligonucleotides targeting PRKCD and two of three targeting GAK caused significant inhibition of DFP-induced mitophagy in the U2OS IMLS cells (Fig. 3a, Supplementary Fig. 3). The level of citrate synthase activity was significantly decreased in DFP treated control cells, this was not the case in cells depleted of PRKCD or GAK, further indicating a reduction in mitophagy upon depletion of these kinases (Fig. 3c). Some loss of citrate synthase was still observed, likely due to incomplete target knockdown as noted by qPCR in the secondary screen (Supplementary Fig. 1a). To understand whether GAK or PRKCD localise to mitochondria upon DFP treatment, U2OS cells were treated or not with DFP and then permeabilised and fractionated by differential centrifugation to enrich for mitochondria. Successful fractionation was confirmed by enrichment of the mitochondrial proteins TIM23 and COX-IV in the mitochondrial fraction (Fig. 3d). PRKCD was strongly enriched in mitochondrial fractions under both control and DFP inducing conditions, while GAK was absent from the mitochondria fraction (Fig. 3d). Mitochondrial localisation of PRKCD was confirmed by immunofluorescence staining of endogenous PRKCD in U2OS IMLS cells, showing a striking co-localisation with the IMLS-EGFP-mCherry reporter both in the presence and absence of DFP (Fig. 3e). Upon induction of mitophagy by DFP, PRKCD was seen with red only structures, indicating it follows mitochondria to the lysosome. Indeed, western blotting confirmed the loss of PRKCD in a DFP-dependent manner that was rescued by BafA1 treatment (Fig. 3f,g). Attempts to identify endogenous GAK localisation by immunofluorescence were not successful with non-specific staining observed with all antibodies tested.

### The kinase activity of GAK and PRKCD are required for functional mitophagy

As GAK and PRKCD are both serine-threonine protein kinases we next sought to determine whether their kinase activities are important for DFP-induced mitophagy. We first tested two recently published kinase inhibitors targeting GAK, IVAP1966 and IVAP1967, with a Kd of 80nM and 190nM respectively (Supplementary Fig. 4a)^21^. Concomitant dosing of IMLS cells with DFP and these GAK inhibitors demonstrated dose-dependent inhibition of mitophagy with IVAP1967, while IVAP1966 had no effect (Supplementary Fig. 4b). Fortunately, we were able to take advantage of a more potent and specific GAK inhibitor (SGC-GAK-1, termed GAKi here) with a Kd of 3.1nM and IC_50_ of 110nM, which also has a negative control probe (SGC-GAK-1N, termed GAKc here, GAK IC_50_ = >50μM) (Supplementary Fig. 4a) as well as a second control that accounts for off-target effects of GAKi (HY-19764, RIPK2 inhibitor)^22^. Using this inhibitor set, we found that DFP-induced mitophagy was significantly reduced in a dose-dependent manner with GAKi by 40% and 60% at 5 μM and 10 μM, respectively (Fig. 4a,b). By contrast, neither GAKc nor RIPK2i had an inhibitory effect (Fig. 4a,b), providing evidence that GAK kinase activity is required for functional DFP mitophagy. The effect of GAKi on mitophagy was not due to reduced cell growth during DFP treatment (Supplementary Fig. 4c).

**Figure 4.**
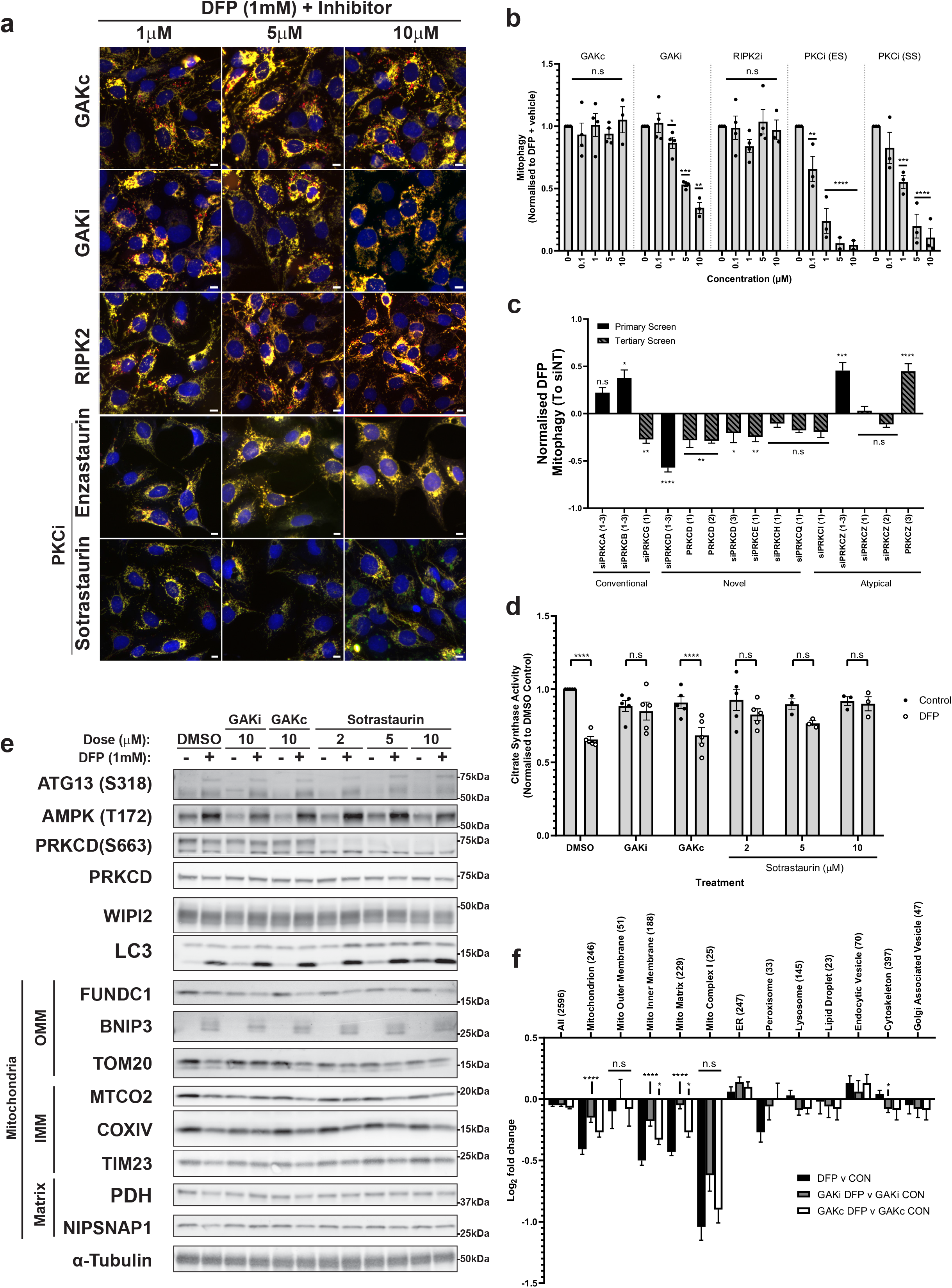
GAK and PRKCD kinase activity regulates DFP-induced mitophagy. **a** Fluorescence images of U2OS IMLS cells treated with 1 mM DFP for 24 h in the presence of indicated inhibitors at stated concentrations (1-10 μM). Scale bar = 10 μm **b** Quantitation of cells red only structures normalised to DMSO + DFP control treatment. Significance was determined by two-way ANOVA followed by Dunnett’s multiple comparison test to the DMSO + DFP control from a minimum of n=3 independent experiments. **c** U2OS IMLS cells treated with siRNA against indicated PKC isoforms (Primary screen = 7.5 nM, Tertiary = 15 nM) for 48 h prior to induction of mitophagy with 1 mM DFP for 24 h. Value in brackets represents oligonucleotide # used. Values represent mean fold change in mitophagy relative to the siNT control ± SEM from n=3 independent experiments. Significance was determined by one-way ANOVA to the relevant siNT control. **d** Citrate synthase activity of U2OS cells treated 24 h ± 1 mM DFP in combination with 10 μM GAKi, GAKc or 2-10 μM Sotrastaurin. Values represent mean citrate synthase activity normalised to DMSO control and plotted ± SEM from n=3 (Sotrastaurin 5/10μm) or n=5 independent experiments. Significance was determined by two-way ANOVA followed by Sidak’s multiple comparison test. **e** U2OS cells were treated ± 1 mM DFP 24 h with GAKi or GAKc (10 μM), Sotrastaurin (2-10 μM) or DMSO control and western blotted for indicated proteins, including outer mitochondrial membrane (OMM), inner mitochondrial membrane (IMM) or Matrix proteins. **f** Cellular protein abundance was determined by mass spectrometry. U2OS cells were treated ± 1 mM DFP for 24 h in addition to DMSO, GAKi or GAKc (both 10 μM) and analysed by mass spectrometry. Comparison of mean abundance of GO annotated proteins between control and DFP treated samples for each GAKi, GAKc and DMSO control are shown ± SEM, value in brackets represent number of proteins classified in group by GO analysis. Significance was determined by two-way ANOVA followed by Dunnett’s post-test to the DFP v CON sample. Significance is denoted in figure where: * = p < 0.05, ** = p < 0.01, *** = p < 0.001, **** = p < 0.0001 and n.s = not significant in all relevant panels.

To investigate a role of the PRKCD kinase activity in mitophagy we utilised the PKC family inhibitors (PKCi) Enzastaurin (ES) and Sotrastaurin (SS), neither of which is solely specific for PKC Delta due to isozyme similarity within the PKC family. Both compounds strongly inhibited DFP-induced mitophagy in a dose-dependent manner (Fig. 4a,b) with Enzastaurin slightly more potent than Sotrastaurin. The conventional (PRKCA, PRKCB), novel (PRKCD) and atypical (PRKCZ) PKC family members were included in the primary siRNA screen, with only siPRKCD showing significantly reduced DFP-induced mitophagy of −57% (Fig. 2c). To further investigate the potential effects of the other PKC isozymes, all PKC isoforms were depleted in the U2OS IMLS cells, which demonstrated that multiple novel PKC isoforms decreased DFP-induced mitophagy (Fig. 4c). Novel PKCs do not require calcium, but diacylglycerol (DAG) for regulation of activity, suggesting this may be important for mitophagy. Use of pan-PKC kinase inhibitors may therefore be beneficial for exploring the role of PKC family members in DFP-induced mitophagy.

To further confirm a role for GAK and PRKCD kinase activities in mitophagy, we determined the level of citrate synthase activity following treatment with GAKi, GAKc and PKCi. As seen in Figure 4d, GAKi and PKCi both strongly blocked DFP-induced loss of citrate synthase whilst this was not seen with DMSO or GAKc control samples. Moreover, addition of GAKi or PKCi both decreased the DFP-induced loss of inner mitochondrial membrane proteins (MTCO2, COXIV, TIM23) and matrix proteins (PDH, NIPSNAP1) as examined by western blot analysis, while loss of the outer mitochondrial proteins FUNDC1 or TOM20 was generally not affected (except that GAKi blocked loss of TOM20) (Fig. 4e). This is perhaps not surprising as some outer mitochondrial proteins are known to undergo proteasomal degradation, such as MFN2, upon induction of DFP-induced mitophagy ^8^. We further confirmed the loss of mitochondrial proteins by analysis of whole-cell protein abundance by mass spectrometry between DMSO and DFP treatments in the presence of GAKi and GAKc. As shown earlier, DFP treatment caused a loss of mitochondrial proteins (determined by GO Analysis), which was reduced for cells treated with GAKi, but not for GAKc, indicating that turnover of mitochondria is hindered when GAK kinase activity is blocked (Fig. 4f).

### GAK and PRKCD do not regulate the HIF1α pathway

To understand how depletion or inhibition of GAK or PRKCD prevents DFP-induced mitophagy, we first examined whether treatment with GAKi or PKCi affected the HIF1α pathway. HIF1α stabilization and expression of the HIF1α responsive genes BNIP3 and BNIP3L (NIX) are important for driving mitophagy triggered by iron chelation with DFP ^8,23^. As expected, BNIP3/3L mRNA and protein levels were induced upon DFP treatment with no change in HIF1α mRNA levels (Fig. 4e and Supplementary Fig. 5a,b). The expression levels of BNIP3/3L were not significantly affected by treatment with GAKi, suggesting that GAK does not regulate mitophagy through the HIF1α response. Furthermore, the PKCi Sotrastaurin had no significant effect on BNIP3/3L expression, although Enzastaurin somewhat reduced the BNIP3L transcript level (Supplementary Fig. 5a), which may explain the relatively higher potency of Enzastaurin blocking mitophagy (Fig. 4b). We focused on Sotrastaurin for further experiments as our data indicated there was an important role for the kinase activity beyond the decrease in BNIP3L seen with Enzastaurin.

BNIP3/3L function as autophagy receptors during HIF1α induced mitophagy by recruiting ATG8 proteins through specific LC3 interacting regions (LIRs) and phosphorylation promotes their function in mitophagy ^24,25^. To check whether BNIP3/3L are targets for GAK or PRKCD mediated phosphorylation, cell lysates from U2OS cells treated or not with DFP together with GAKi or PKCi were analysed using phos-tag acrylamide gels that significantly retard the migration of phosphorylated proteins ^26^. Importantly, neither the protein expression of BNIP3/3L with DFP nor their migration patterns were changed in GAKi, GAKc and PKCi treated samples (Supplementary Fig. 5b), indicating that neither BNIP3 nor BNIP3L are targets for direct phosphorylation by PRKCD or GAK.

### GAK and PRKCD kinase activities do not regulate PRKN-dependent mitophagy or starvation-induced autophagy

As both GAK and PRKCD regulate DFP-induced mitophagy without affecting the HIF1α pathway, we next sought to examine whether these kinases also regulated PRKN-dependent mitophagy. The U2OS IMLS-EGFP-mCherry cell line was transduced with lentivirus to constitutively overexpress PRKN (untagged), which permits strong induction of mitophagy in response to mitochondrial depolarisation such as that induced by the H^+^ ionophore CCCP ^27,28^. Indeed, treatment with CCCP for 16 h led to significant induction of mitophagy and near-total loss of mitochondrial network as seen by accumulation of red only structures (Fig. 5a), reduced citrate synthase activity (Fig. 5b) as well as the loss of the mitochondrial matrix protein PDH, along with PRKCD that also localises on mitochondria (Fig. 5c,d). Importantly, we could block PRKN-dependent CCCP-induced mitophagy by co-treatment with the ULK1/2 inhibitor MRT68921 or the lysosomal inhibitor BafA1 (Fig. 5a-d) ^13,29^. Co-treatment with GAKi, GAKc or PKCi had no effect (Fig. 5a-d), suggesting that GAK and PRKCD kinase activities are dispensable for PRKN-mediated mitophagy under these conditions. We next considered whether GAKi or PKCi could impair starvation-induced autophagy. Cells were incubated in nutrient starvation media (EBSS) for 2 h and the autophagic flux examined by immunostaining for endogenous LC3B in the absence or presence of BafA1, as analysed by fluorescence microscopy (Fig. 5e,f) or immunoblotting (Fig. 5g). Whilst incubation in EBSS increased LC3 flux relative to the control, addition of GAKi or GAKc to EBSS treated cells did not significantly inhibit the autophagic flux. PKCi caused elevated LC3-II levels in the absence of BafA1.

**Figure 5.**
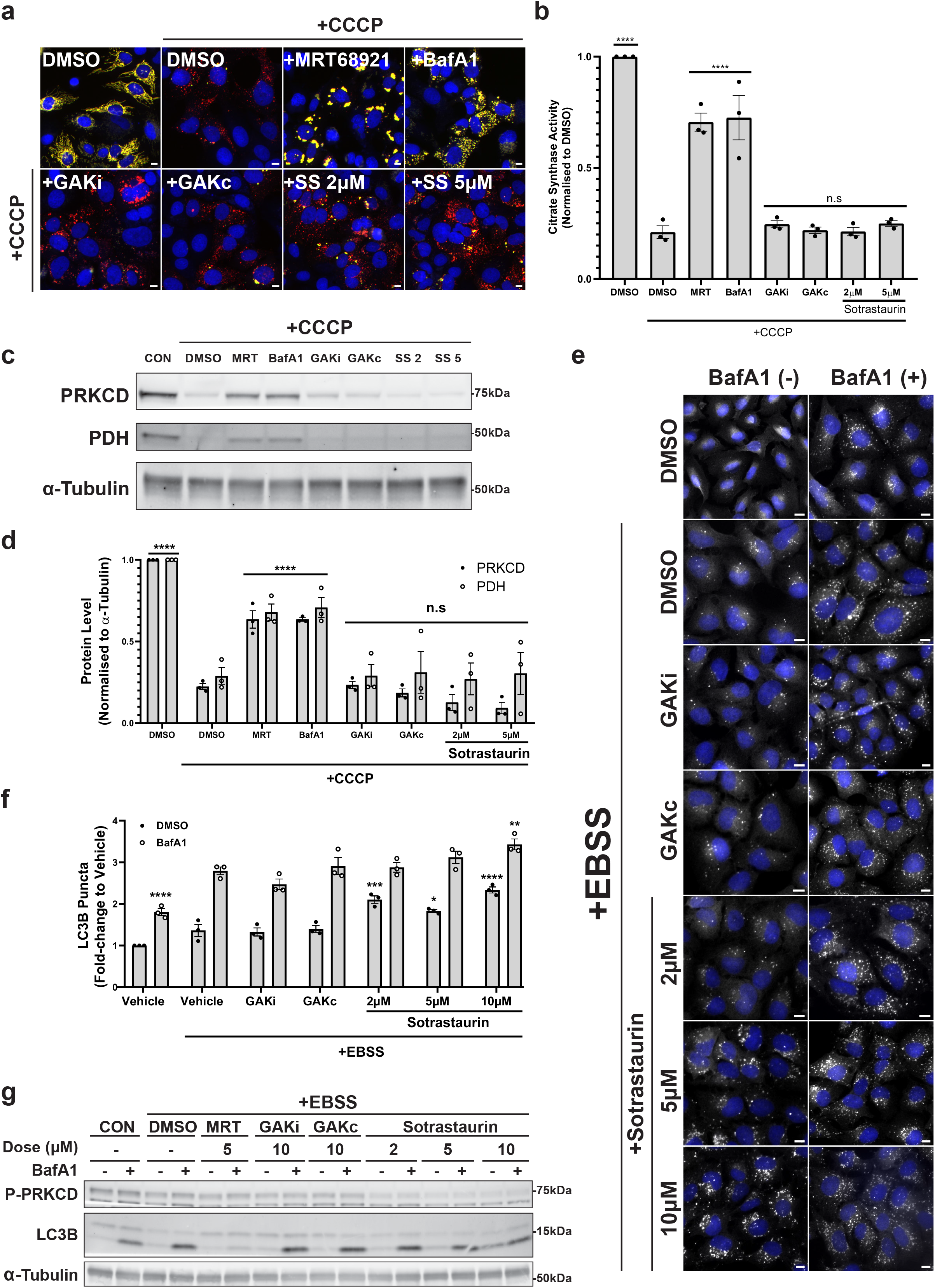
GAK and PRKCD kinase activity are dispensable for PRKN-dependent mitophagy. **a** Representative fluorescence images of U2OS IMLS-PRKN cells treated for 16 h ± 20 μM CCCP and including 10 μM QVD-OPh to promote cell survival in addition to either MRT68921 (5 μM), BafA1 (50 nM), GAKi (10 μM), GAKc (10 μM) or Sotrastaurin (2-5 μM). Scale bar = 10μm **b** Cells treated as in **a** and assayed for citrate synthase activity and normalised to DMSO control. Mean value plotted ± SEM from n=3 independent experiments and significance determined by one-way ANOVA followed by Dunnett’s multiple comparison to the CCCP+DMSO control. **c** Representative example of western blots from cells treated as in **a** and blotted for indicated proteins. **d** Quantitation of PRKCD and PDH levels from western blots in **c** from n=3 independent experiments ± SEM. Values represent protein level normalised first to α-Tubulin and subsequently normalised to the DMSO control. Significance was determined by two-way ANOVA followed by Dunnett’s multiple comparison test to the DMSO control. **e** Representative 20x immunofluorescence images of U2OS cells stained for endogenous LC3B and nuclei (DAPI, blue). Cells were grown in complete media or EBSS (starvation) media for 2 h with addition of GAKi (10 μM), GAKc (10 μM) or Sotrastaurin (2-10μM) ± 50 nM BafA1, scale bar = 10 μm. **f** Quantitation of LC3B puncta from **e**. The average LC3 puncta per cell was normalised to that of the complete media control and represents the mean ± SEM from n=3 independent experiments. Significance was determined by two-way ANOVA followed by Dunnett’s multiple comparison test to the EBSS vehicle treated sample. **g** Representative western blot of cells treated as in **e** and blotted for indicated proteins. * = p < 0.05, ** = p < 0.01, *** = p < 0.001 and **** = p < 0.0001 and n.s = not significant in all relevant panels.

Taken together, we show that while GAK and PRKCD kinase activities are required for efficient DFP-induced mitophagy, but are dispensable for PRKN-dependent mitophagy and starvation-induced autophagy.

### PRKCD inhibitors reduce ULK1 initiation complex assembly

To further elucidate how GAK or PRKCD kinase inhibition may block DFP-induced mitophagy, we examined the recruitment of core autophagy machinery components to mitochondria. U2OS IMLS cells treated with DMSO or DFP were immunostained for endogenous ATG13, LC3B, ULK1 or WIPI2 and examined by confocal microscopy. In many cases, the early autophagosome structures induced by DFP treatment were localised at or close to mitochondria, likely representing mitophagosome start sites (Fig. 6a-d). Using high-content imaging we found that formation of WIPI2, ATG13 or ULK1 puncta were all strongly induced (~5-fold) in response to 24h DFP treatment (Fig. 6e,f, Supplementary Fig. 6a-d). Treatment with GAKi or GAKc did not affect the number of WIPI2, ATG13 or ULK1 puncta observed following 24 h of DFP treatment (Fig. 6e,f, Supplementary Fig. 6a-d). Moreover, neither GAKi nor GAKc affected phosphorylation of ULK1 at S555 (AMPK site) or S757 (mTOR site) (Supplementary Fig. 6e), suggesting that GAK regulates DFP-induced mitophagy through a different mechanism. Interestingly, control cells treated with GAKi only (no DFP) displayed an increased number of WIPI2 puncta (Supplementary Fig. 6a,d), but the reason for this is not apparent.

**Figure 6.**
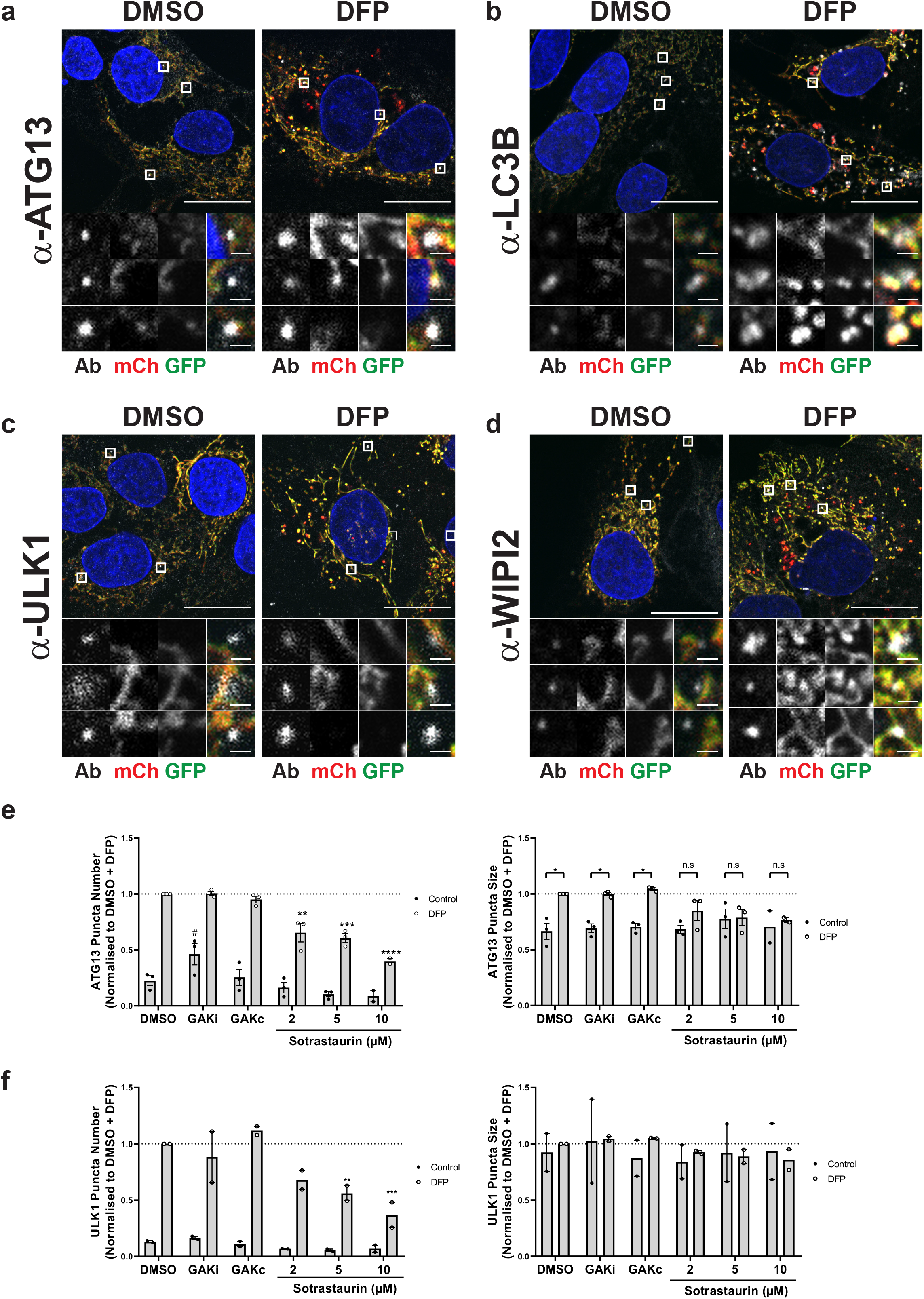
Early autophagy protein recruitment is defective upon PKCi. **a-d** U2OS IMLS cells were treated ± 1 mM DFP for 24 h, fixed and stained for nuclei (DAPI) and the indicated endogenous autophagy markers; **a** ATG13, **b** LC3B, **c** ULK1 or **d** WIPI2. Representative 63x images of cells taken by Zeiss LSM 710 are shown, scale bar = 10 μm. **e-f** U2OS cells were treated ± 1 mM DFP for 24 h together with GAKi (10 μM), GAKc (10 μM) or Sotrastaurin (2-10 μM), then fixed in PFA before staining for nuclei (DAPI) and the indicated endogenous early autophagy markers; **e** ATG13, **f** ULK1. The number and size of puncta formed for each marker was analysed and values obtained were normalised to the DMSO + DFP control. Mean values were plotted from n=3 independent experiments ± SEM. Significance was determined by two-way ANOVA and Dunnett’s multiple comparisons test to the DMSO+DFP control sample where * = p < 0.05, ** = p < 0.01, *** = p < 0.001, **** = p < 0.0001, n.s = not significant and # = p < 0.05 (to the DMSO Control).

Most importantly, co-treatment of cells with DFP and PKCi caused a dose-responsive decrease in the number and size of ULK1 and ATG13 puncta (and to a lesser extent WIPI2 though this was not significant) (Fig. 6e,f and Supplementary Fig. 7d). However, neither the phosphorylation of ATG13 at S318 (a known ULK1 site) ^30^ or the phosphorylation of AMPK at T172 (an activation site) ^31^ that occur in response to DFP treatment were impaired or altered by co-treatment with GAKi, GAKc or PKCi (Fig. 4e). We thus conclude that the kinase activity of PRKCD is required for successful assembly and formation of the ULK1 complex during DFP-induced mitophagy, but that it does not affect ULK1 activity directly. As PRKCD is localised to mitochondria, a failure to generate an initiation structure upon its depletion or inactivation likely explains the reduced level of mitophagy observed.

### GAK inhibition alters mitochondrial morphology

As GAK inhibition did not affect HIF1α signalling, ULK1 activity or ATG13/ULK1 recruitment, we next examined mitochondrial morphology. IMLS cells co-treated with DFP and siGAK (Supplementary Fig. 3) or GAKi (Fig. 4a) demonstrated abnormal mitochondrial network structures compared to control cells. Using live-cell microscopy, we observed a collapsed mitochondrial network with clumped mitochondrial regions and reduced red structures in IMLS cells co-treated with DFP and GAKi that were not seen with GAKc (Supplementary Movie 1). Confocal microscopy confirmed this and by contrast, cells co-treated with DFP and PKCi retained an elongated network (Fig. 7a). Cells treated with a combination of Oligomycin and Antimycin A (O+A) or CCCP, known to depolarize mitochondria, caused fragmented network phenotypes (Fig. 7a). In the latter case, little mitophagy is seen in response to mitochondrial depolarisation as this was carried out in IMLS cells without PRKN overexpression. To quantify mitochondria morphology, images of U2OS cells depleted of the fission enzyme DRP1 or the fusion enzyme OPA1 (causing tubulated or fragmented mitochondrial phenotypes respectively, Supplementary Fig. 7), were applied to train a cell classifier in CellProfiler Analyst. Applying this, we confirmed that GAKi treated cells exhibit a hyper fused mitochondria phenotype while no morphological differences were detected in GAKc or PKCi treated samples (Fig. 7b). The observed GAKi phenotype was not caused by a general defect in the fission/fusion rate of the mitochondrial network, as mitochondrial fragmentation could still be induced by co-treatment of GAKi with DFP and CCCP (Fig. 7c). Importantly, GAKi still prevented the formation of red only structures under such conditions, indicating that GAK is important for proper uptake of fragmented mitochondria into autophagosomes.

**Figure 7.**
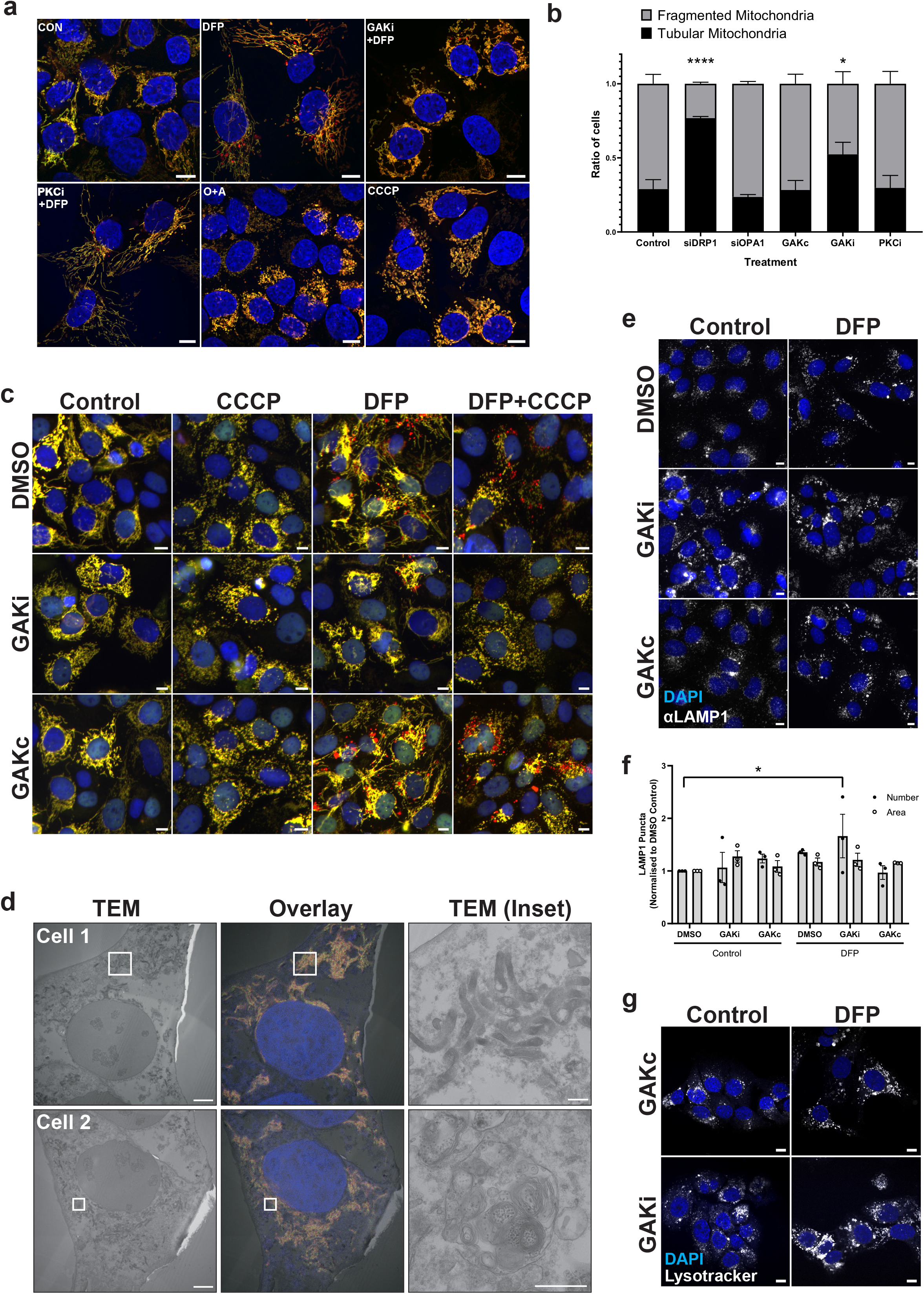
GAKi induces abnormal mitochondrial and lysosomal morphology. **a** Representative 63x images of U2OS IMLS cells taken by Zeiss LSM 710 confocal microscopy. Cells were treated ± 1 mM DFP 24 h in addition to GAKi (10 μM), Sotrastaurin (PKCi – 2 μM), Oligomycin and Antimycin A (O+A – 10 μM and 1 μM respectively) or CCCP (20 μM), scale bar = 10 μm. **b** Machine learning classification of U2OS IMLS cell mitochondrial network as fragmented or tubular (utilising EGFP images, see methods) after 24 h treatment with GAKi (10 μM), GAKc (10 μM) or Sotrastaurin (PKCi – 2μM) compared to 72 h knockdown of non-targeting control, siDRP1 or siOPA1. Significance was determined by two-way ANOVA followed by Dunnett’s post-test to the control treatment. **c** U2OS IMLS cells were treated as indicated with DMSO, GAKi or GAKc (10 μM each) for 24 h in addition to either DFP (24h, 1 mM), CCCP (20 μM, 12 h) or in combination. Images obtained by 20x objective, scale bar = 10μm. **d** U2OS IMLS cells were treated with 1 mM DFP + 10 μM GAKi for 24h prior to fixation for CLEM analysis. EM images demonstrate mitochondrial clustering (Cell 1) and an increase in autolysosome structures (Cell 2) induced by GAKi treatment, scale bar = 10 μm, inset = 1 μM. **e** U2OS cells treated ± 1 mM DFP 24 h in addition to DMSO, GAKi (10μM) or GAKc (10μM) were PFA fixed and subsequently stained for endogenous LAMP1. Images acquired by widefield microscopy on a Zeiss AxioObserver microscope, scale bar=10 μm. **f** Quantitation of LAMP1 structures identified in **e** for size and number from n=3 independent experiments and plotted as mean ± SEM. Significance was determined by two-way ANOVA followed by Dunnett’s multiple comparisons test to the DMSO control. **g** U2OS cells were treated for 24 h ± 1 mM DFP with either GAKc (10 μM) or GAKi (10 μM) and then stained for lysosomes using lysotracker red DND-99 at 50 nM. Representative images taken by Zeiss LSM710, scale bar = 10 μm. Significance was denoted where * = p < 0.05, **** = p < 0.0001 and n.s = not significant.

We next examined GAKi treated U2OS IMLS cells by CLEM. Interestingly, this showed condensed parallel layers of mitochondria that were not fused, but rather stacked closely with one another (Fig. 7d). Additionally, large autolysosome structures were observed, which may indicate lysosomal defects following GAKi treatment (Fig. 7d). Indeed, an increase in the number of lysosomal structures was detected in cells stained for the late endosome/lysosome marker LAMP1 following co-treatment with DFP and GAKi (Fig. 7e,f). As GAKi did not inhibit PRKN-dependent mitophagy (Fig. 5a-d) or starvation-induced autophagy (Fig. 5e-g) we reasoned that the large autolysosomes seen in GAKi treated cells retain their acidity and degradative capacity. In line with this, lysotracker positive structures were detected in cells treated with GAKi or GAKc in the presence or absence of DFP (Fig. 7g). Thus, we believe that the mitophagy defect induced by GAKi is due to inefficient cargo loading or delivery to lysosomes rather than a lysosomal defect.

### Mass Spectrometry with GAKi

To try to identify relevant substrates for GAK kinase-dependent regulation of DFP-induced mitophagy, we carried out phospho-proteomic analysis of cells treated with DFP in combination with GAKi or GAKc. GAK is currently defined as an understudied kinase ^32^ and GAKi was developed by the SGC consortium to reveal novel understandings of GAK cellular function ^22^. Analysis of both the protein abundance and phospho-sites of interest (Fig. 8a) demonstrated that DFP treatment increased the abundance of several proteins in a GAKi independent manner, including HIF1α, BNIP3L, Hexokinase I/II, LAMP1/2 and LC3B. In contrast, SQSTM1 abundance decreased in response to DFP even with GAKi, further showing that lysosomes are functional in this state. Interestingly, phosphorylation of RAB7A at S72 was observed in response to DFP treatment. The same RAB7A modification was recently reported to facilitate a key step in PRKN-dependent mitophagy, suggesting that similarities exist during DFP mitophagy ^33^. RAB14 was also phosphorylated (at S180) in response to DFP, a change that may be interesting to examine further.

**Figure 8.**
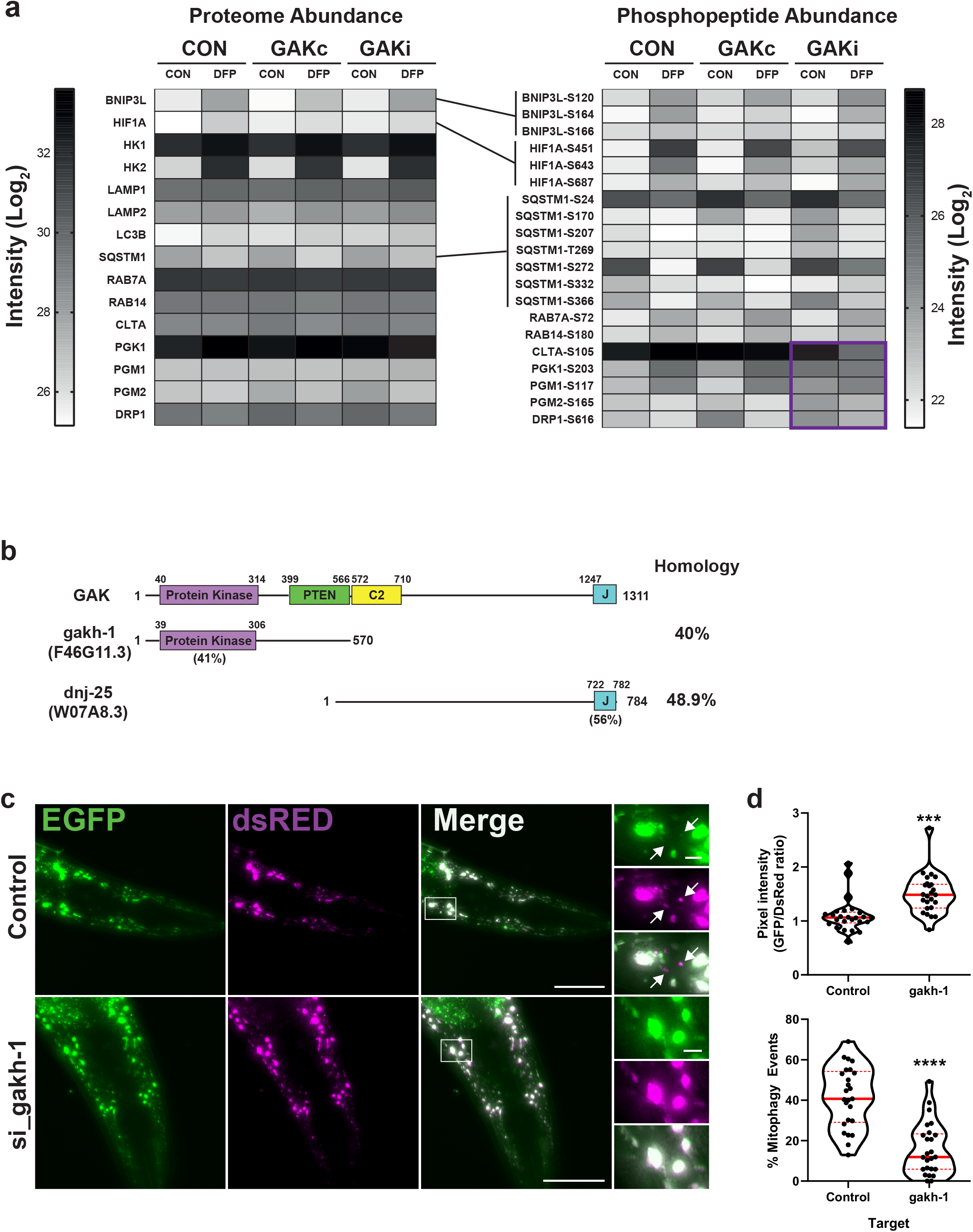
gakh-1 regulates mitophagy in vivo. **a** Mass spectrometry of DFP and GAKi regulated proteins/phospho-peptides. U2OS cells were treated ± 1 mM DFP for 24 h in combination with DMSO vehicle, GAKi or GAKc (both 10 μM). Cell pellets were collected and split between proteome and phospho-peptide analysis (see methods). Data represents average intensity from n=2 independent experiments. **b** Schematic representation of GAK domain structure and orthologues gakh-1 and dnj-25 present in C.elegans. Homology values were obtained by protein blast alignment. **c** In vivo detection of mitophagy in C. elegans. Transgenic nematodes expressing mtRosella in bodywall muscle cells were treated with gakh-1 RNAi or pL4440 control vector. dsRED only structures represent mitochondria in acidic compartments (arrowheads). Scale bar = 50 μm, inset = 5 μm. **d** Mitophagy stimulation signified by the ratio between pH-sensitive GFP to pH insensitive dsREd (n= 25, upper panel). Quantification of the frequency of mitochondria undergoing mitophagy (dsREd puncta lacking EGFP co-localisation) are expressed as percentage of total mitochondria detected (n= 25, lower panel). The data is presented as violin plots of individual values with median (red, solid line) and quartiles (red, dashed line) shown. Significance was determined by unpaired two tailed t-test from n=2 independent experiments, where *** = p < 0.001 or **** = p < 0.0001.

GAKi-dependent changes in phosphorylation were also seen, we observed a very specific loss of clathrin light chain A (CLTA) phosphorylation at S105. This site lies at the interface between the clathrin heavy and light chains, supporting a previously described role of GAK in clathrin-cage uncoating. Increased levels of PGK1 S203, PGM1 S117 and PGM2 S165 (established phosphorylation sites to induce glycolysis) were detected upon GAKi treatment without DFP, suggesting that glycolysis may be higher at the basal state. We also see higher DRP1 S616 levels with GAKi and DFP, a modification normally associated with increased fission activity that can be mediated by CDK1 ^34^. Moreover, SQSTM1 phosphorylation at S272 is increased with GAKi, also previously noted to be mediated by CDK1 ^35^, suggesting that GAKi treatment increases CDK1 activity and may indicate increased cell numbers in mitosis ^36^.

### GAK and PRKCD modulate mitophagy in vivo

Knockout of GAK orthologues have been shown to cause embryonic lethality in mice, *C.elegans* (*dnj-25*) and *D.melanogaster* (*dAux*), which is postulated to occur due to defective clathrin-dependent endocytosis, as uncoating of clathrin-coated vesicles is mediated by the J-domain of GAK ^37–39^. Indeed, expression of the J-domain alone is able to rescue survival in both mice and drosophila GAK knockout models ^39,40^. *C.elegans* contains two orthologues of GAK, comprising the kinase domain (gakh-1, F46G11.3, 40% overall homology) or the J-domain (dnj-25, W07A8.3, 49% overall homology) (Fig. 8b), where targeting of the latter by RNAi causes lethality during larval development ^38^. As our data indicate a requirement of GAK kinase activity for efficient mitophagy in mammalian cells, we examined the effect of targeting gakh-1 upon basal mitophagy in a *C.elegans* reporter line expressing mtRosella GFP-DsRed in body wall muscle cells, following the concept of the IMLS reporter cell lines. *C.elegans* fed gakh-1 RNAi demonstrated a significant decrease in the ratio of GFP to DsRed and more mitophagy events (DSRed only structures, Fig. 8c) compared to the RNAi control, confirming that GAK kinase activity is important for basal mitophagy *in vivo*.

We next sought to examine the role of PRKCD in mitophagy *in vivo* and targeted Prkcd in our recently established transgenic mitophagy reporter zebrafish line, expressing zebrafish Cox8 fused to EGFP-mCherry (Abudu et al., 2019). Zebrafish contain paralogues (prkcda and prkcdb) that are ~80% similar to one another and ~76% homologous to human PRKCD, with all major domains being highly conserved (Fig. 9a). We examined the spatio-temporal expression pattern of *prkcda* and *prkcdb* during zebrafish development up until 5 days post-fertilisation (dpf). Whilst a high level of maternal *prkcd*a mRNA was observed at 2 hours post fertilisation (hpf), possibly indicating a role in early embryonic signalling events, both genes were expressed at similar levels throughout development (Fig. 9b). The spatial distribution of *prkcda* and *prkcdb* were analysed by whole-mount *in situ* hybridisation (WM-ISH) at 5 dpf compared to a sense probe negative control and showed strong staining in the corpus cerebelli region of the hindbrain for both genes (Fig. 9c, Supplementary Fig. 8a), this is in agreement with ISH data deposited in the ZFIN database (http://zfin.org). *prkcda* expression was also detected in the eyes and the olfactory bulbs (Fig. 9c) whereas *prkcdb* was found in patches of the retina, optical tectum and the spinal cord (Fig. 9c). We also analysed the spatio-temporal pattern of *gak* expression across zebrafish development and observed that even though it is expressed consistently throughout development (Supplementary Fig. 8b), its spatial expression pattern varies across development with staining of the caudal hindbrain and retina at 2 dpf, the optical tectum, neurocranium, retina and kidney at 3dpf, the kidney and neurocranium at 4dpf and the liver, olfactory bulb, optical tectum and retina at 5dpf of the wild type zebrafish larvae, with no staining of the control probe (Supplementary Fig. 8c).

**Figure 9.**
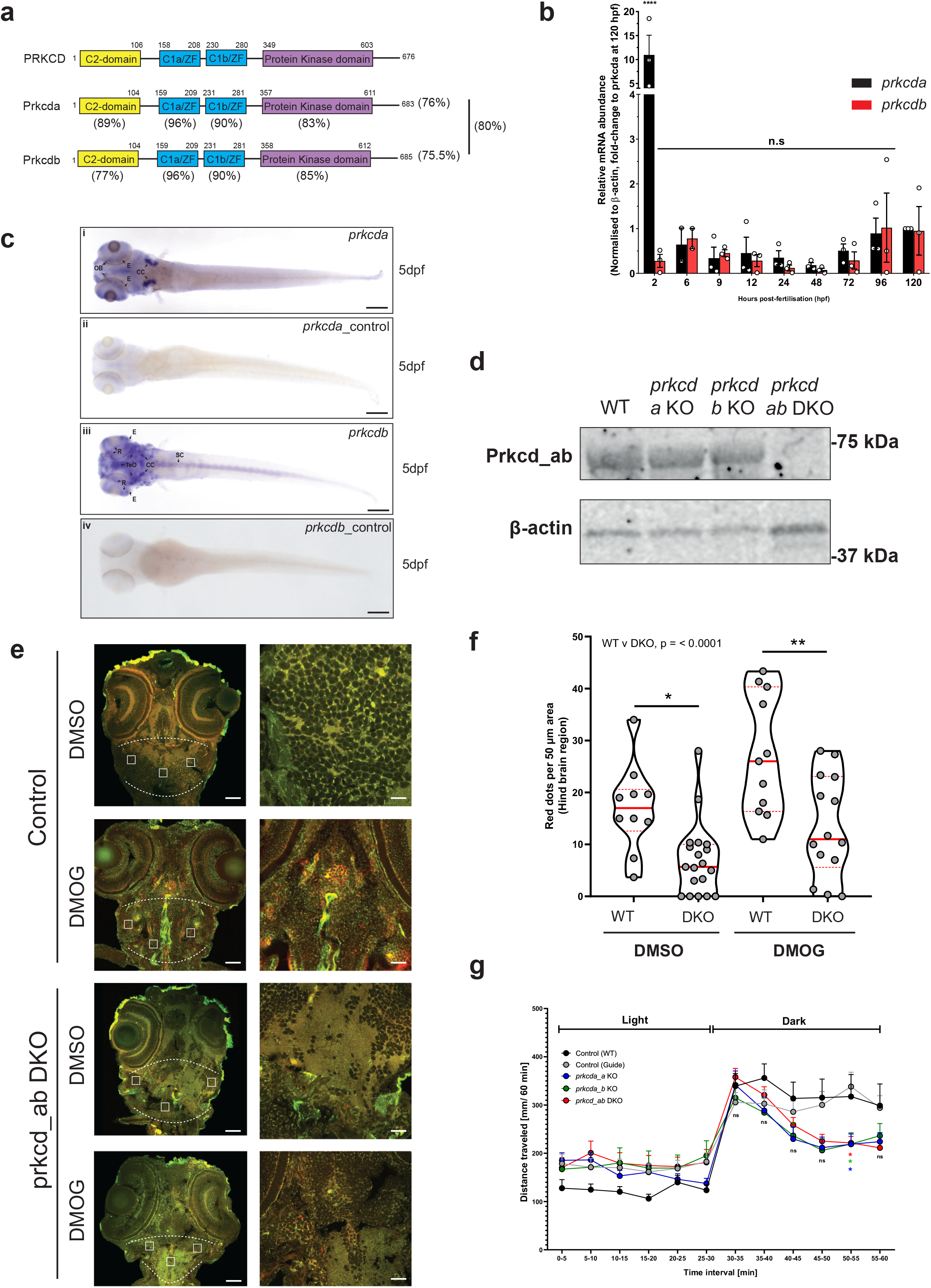
prkcda and prkcdb regulate mitophagy in vivo. **a** Overview and schematic diagram of human PRKCD, zebrafish prkcda and prkcdb proteins. Percentage identity of respective domains on comparison with human counterpart shown below the zebrafish domains. Also shown is the percentage identity of the protein amongst each other. **b** Temporal expression pattern of prkcda and prkcdb. The graph shows the mean relative transcript abundance in whole zebrafish embryos from 2 hpf to 5 dpf from n=2 (6 hpf) or n=3 (all others) independent experiments **c** Spatial expression pattern of prkcda and prkcdb at 5 dpf as demonstrated by whole mount in situ hybridisation at the indicated stage using a 5’UTR probe. Both the larvae are in dorsal view. Scale bar = 200 μm. **d** Representative immunoblots of Prkcd and β-actin on whole embryo lysates from wild-type and single or double prkcda/prkcdb KO (DKO) animals. **e** Representative confocal images of cryosections taken from control (guide only) and prkcd_ab DKO transgenic tandem-tagged mitofish larvae treated with DMSO only or with DMOG at 3 dpf. Images are from the hind brain region of the respective larvae as marked. Scale bars = 50 μm (left), 20 μm (right). **f** The graph shows the average number of red puncta from three 50 μm hind brain region (as marked in e) from each of 10-19 larvae for control and prkcd_ab DKO tandem-tagged mitofish larvae. Significance was determined by two-way ANOVA followed by Tukey’s post-test to compare all groups. **g** Motility analysis of zebrafish embryos at 5 dpf using the “Zebrabox” automated videotracker (Viewpoint). Assay was carried out during daytime, and consisted of one cycle of 30 min exposure to light followed by 30 min of darkness. Data represents mean distance moved ± SEM. Each group consisted of 43-124 larvae from n= 9 independent experiments. Significance was determined by two-way ANOVA followed by Dunnett’s post-test to the Control (WT). Significance are denoted where *p < 0.05, **p < 0.01, ***p < 0.001, **** p < 0.0001, n.s = not significant.

To investigate a possible role for Prkcd in the mitophagy reporter line, we employed CRISPR/Cas9-mediated genome editing in zebrafish embryos using guide sequences targeting *prkcda* and *prkcdb* individually or together (Fig. 9d and Supplementary Fig. 8d), with the latter referred to as *prkcd_ab* double knock-out (DKO). We verified loss of Prkcd protein levels in DKO embryos by western blot (Fig. 9d) and therefore used these embryos for *in vivo* mitophagy analysis.

To induce mitophagy, zebrafish larvae at 2 dpf were incubated with varying concentrations of DFP and DMOG for 24 hours. DFP failed to induce mitophagy at all concentrations, however, DMOG induced mitophagy as shown by reduced levels of Tim23 (Supplementary Fig. 8e,f), in line with a recent report ^41^. We used 100 μM DMOG for further experiments, as significant lethality and morphological defects were present at higher (250 μM) concentrations (Supplementary Fig. 8g). As *prkcd_a/b* express highly in the hindbrain (Fig. 9c), we examined mitophagy in this region of DKO larvae compared to control larvae, both at basal (DMSO) and DMOG-treated conditions. Sections of fixed larva were imaged by confocal microscopy and the number of red puncta in the hindbrain region quantified. The number of red only puncta was significantly reduced in the hindbrain of DKO larvae with both basal and DMOG-induced conditions (Fig. 9e,f), showing an important role of Prkcd_a/b in regulating mitophagy. Interestingly, we noticed the lack of a properly formed eye-lens in one of the two eyes of the *prkcd_ab* DKO, with considerably reduced mitophagy levels when compared to the control (Supplementary Fig. 8h). We also observed an inconsistent and varied movement pattern of the DKO larvae, and therefore quantified their locomotor activity. Tracking and quantification of locomotion was performed in alternating light and dark conditions with a reduced swimming trend in the dark phase for both Prkcd single KO and DKO larvae compared the WT control (Fig. 9g). It is likely that this may be related to the observed lens defect or a hindbrain clustered motor-neuron dependent phenotype.

In conclusion, we show that the activity of both GAK and PRKCD is important for regulation of basal mitophagy *in vivo*, highlighting the evolutionary relevance of these kinases in mitophagy.

## DISCUSSION

In this manuscript, we have tested a panel of putative lipid-binding proteins for their ability to regulate DFP-induced mitophagy. We have identified eleven candidate proteins that demonstrate significant modulation of DFP-induced mitophagy and explored two of these, GAK and PRKCD, in further detail. In both cases, we show that functional kinase activity is required for their positive regulation of DFP-induced mitophagy and we have identified putative mechanisms for how each function in mitophagy regulation. Importantly, neither GAK nor PRKCD are required for PRKN-dependent mitophagy or starvation-induced autophagy, offering novel targets for the specific modulation of PRKN-independent mitophagy. Critically, we find that both of these targets identified *in vitro* are also relevant targets *in vivo* for the regulation of basal mitophagy in *C.elegans* and *D.rario*, demonstrating their conserved function in mitophagy.

The linkage of GAK to mitophagy is of particular relevance to neurodegenerative disease as SNPs in GAK have previously been identified as a risk factor for familial PD ^19^ and expression changes in GAK are observed in the substantia nigra of PD patients ^42^. Knockout of the drosophila homologue of GAK (Auxilin) also demonstrates Parkinsonian like mobility defects and loss of dopaminergic neurons ^43^. Further work has shown that the commonly mutated PD gene, LRRK2, can phosphorylate Auxilin^44^. Studies of GAK have to date, however, been hindered by the limited availability of chemical tools alongside the inherent difficulty of modulating embryonic lethal genes to dissect their function. *C.elegans* presents a unique opportunity for the *in vivo* study of GAK kinase function due to the presence of two orthologues of human GAK, with the homologous kinase domain within a separate protein (gakh-1) than that of the developmentally essential J-domain (dnj-25) ^38^. By knockdown of gakh-1, we were able to see >50% reduction in basal mitophagy in muscle-cell wall highlighting the importance of GAK for mitophagy. Further investigation is required to mechanistically ascertain how this functional effect is mediated, as Hif1α induction and subsequent recruitment of ULK1/ATG13 to mitophagosomes in response to DFP stimulation appears normal in cells with GAK kinase activity inhibited. However, we observed a significant disruption of mitochondrial network morphology which may modulate the ability of mitochondria to be loaded correctly into mitophagosomes. Artificially inducing mitochondrial fragmentation with CCCP, however, was not sufficient to rescue mitochondrial degradation. Moreover, enlarged lysosomes were observed, though these still possess degradative potential to mediate starvation-induced or PRKN-dependent mitophagy. Mass spectrometry analysis indicates that several glycolytic enzymes are activated in response to GAKi treatment, which may indicate that more fundamental metabolic changes are occurring that alter the cellular degradation of mitochondria.

PRKCD is a member of the large PKC kinase family and belongs to the subgroup of novel PKCs (nPKCs) including PRKCE, PRKCH and PRKCQ that are all activated with DAG independent of Ca^2+^. Whilst we have primarily focused upon PRKCD due to its prominent mitochondrial localisation, it is interesting to note that all nPKCs reduced DFP-induced mitophagy to varying extents, suggesting DAG and PKCs play a prominent role in the regulation of mitophagy. Usage of a pan-PKC inhibitor consistently gave a stronger and more robust inhibition of DFP-induced mitophagy, indicating that several isoforms could play a role or serve redundant functions with one another. Treatment with the PKC inhibitor sotrastaurin led to a significant inhibition in the recruitment of early autophagy markers, thereby reducing mitophagosome formation.

Formation of mitochondrial DAG has previously been observed during oxidative stress induced by H_2_O_2_, which resulted in the recruitment of Protein Kinase D1 (PKD1) to mitochondria^45^. Recent data in mice have also implicated PKD1 in the regulation of mitochondrial depolarisation and is itself regulated by PRKCD in the activation loop at S738/S742, it is tempting to speculate that PKD1 might also be involved in DFP-induced mitophagy ^46^. Another intriguing detail is that PKD1 can bind to AKAP13, another identified regulator of mitophagy in our screen ^47^. The formation of mitochondrial DAG could therefore be a key step in the regulation of DFP-dependent mitophagy. DAG can be formed from phosphatidic acid by phosphatidic acid phosphohydrolases, such as the Lipin family of proteins. Interestingly, Lipin-1 deficiency is associated with accumulation of mitochondria combined with morphological abnormalities ^48^. Additionally, Lipin-1 depletion was found to reduce the level of PKD1 phosphorylation with a subsequent decrease in VPS34 mediated PtdIns(3)P formation ^48^. Further examination of PKD1 regulation may therefore be important for studying the role of PKCs in DFP-induced mitophagy.

Interestingly, neither GAK nor PRKCD inhibition was able to modulate PRKN-mediated mitophagy. This adds further evidence that the machinery required for PRKN-dependent and independent mitophagy pathways are fundamentally different. Tantalisingly, both *c.elegans* and *d.rario* models indicate that basal mitophagy (along with DMOG-stimulated in zebrafish experiments) can be regulated by these kinases as opposed to stress-induced mitophagy, such as that regulated by PRKN. By looking at the hindbrain region of zebrafish, showing the highest expression of Prkcd, we observed a significant reduction in the level of mitophagy upon depletion of prkcda and prkcdb and impairment of locomotory responses. This may be due to deterioration of hindbrain locomotory neurons as a consequence of impaired mitophagy or may be associated with an anxiety-like response (thought to be triggered in zebrafish due to light changes) noted previously in mice to be impaired with PRKCD depletion^49^.

To conclude, this initial screen of lipid-binding proteins in DFP-induced mitophagy identified two lipid-binding kinases that have been validated by functional characterisation and confirmation in higher organisms. This highlights the importance of protein-lipid interactions and provides a strong initial basis for further investigation into the molecular mechanisms of mitophagy.

## Materials and Methods

### Materials

Lysotracker Red DND-99 (L7528) was from ThermoFisher Scientific. Antimycin A (A8674), DFP (379409), DMOG (D3695), SGC-GAK-1 (GAKi, SML2202), SGC-GAK-1N (GAKc, SML2203) and Q-VD-OPh (SML0063) were from Sigma Aldrich. Bafilomycin A1 (BML-CM110), CCCP (BML-CM124) were from Enzo Life Sciences. Enzastaurin (S1055), Oligomycin A (S1478), and Sotrastaurin (S2791) were from Selleckchem. MRT68921 (1190379-70-4) and VPS34-IN1 (1383716-33-3) were from Cayman Chemical. IVAP1966 (12g) and IVAP1966 (12i) were gratefully received from the lab of Prof. Piet Herdewijn ^21^. HY-19764 was gratefully received from the structural genomics consortium ^22^. Bradford reagent dye (#5000006) was from Bio-Rad. 1,4-dithiothreitol (DTT, #441496P) was from VWR. Complete EDTA-free protease inhibitors (#05056489001) and phosphatase inhibitors (#04906837001) were from Roche.

### Cell Lines, Maintenance and Induction of Mitophagy

U2OS FlpIN TRex cells with stable dox-inducible expression of MLS-EGFP-mCherry (referred to as IMLS cells) ^11^ were grown and maintained in a complete medium of Dulbecco’s Modified Eagle Medium (DMEM – Lonza 12-741F) supplemented with 10% v/v foetal bovine serum (FBS – Sigma Aldrich #F7524) and 100 U/ml Penicillin + 100 μg/ml Streptomycin (ThermoFisher Scientific #15140122) in a humidified incubator at 37°C with 5% CO_2_. U2OS IMLS cells with stable expression of PRKN were generated by cloning of PRKN into a pLenti-III-PGK viral expression vector that was co-transfected into 293FT cells with psPAX2 and pCMV-VSVG to generate lentiviral particles, which were transduced into U2OS IMLS cells and positive cells selected with puromycin (Sigma #P7255).

Mitophagy was typically induced utilising 1 mM DFP by addition to cell culture media for 24 h. In the case of PRKN overexpression, CCCP was used at 20 μM for 16 h or a combination of Oligomycin and Antimycin A (10μM and 1μM respectively) for 16 h. In the case of PRKN-dependent mitophagy experiments, the pan-caspase inhibitor Q-VD-OPh ^50^ was included to reduce cell death and improve imaging quality, in accordance with previous papers studying PRKN-dependent mitophagy ^51^.

### Imaging and Image Analysis

The initial siRNA screen, secondary siRNA screen and other experiments where indicated were carried out utilising an AxioObserver widefield microscope (Zen Blue 2.3, Zeiss) with a 20x objective (NA 0.5). Relevant channels were imaged using a solid-state light source (Colibri 7) and multi-bandpass filter (BP425/30, 524/50, 688/145) or individual filters. The tertiary siRNA screen was carried out utilising an ImageXpress Micro Confocal (Molecular Devices) using a 20x objective (NA 0.45). Confocal images were taken with a LSM710 microscope or LSM800 (Zebrafish experiments) microscope (Zen Black 2012 SP5 FP3, Zeiss) utilising a 63x oil objective (NA 1.4) combined with a laser diode (405nm), Ar-Laser Multiline (458/488/514nm), DPSS (561nm) and HeNe-laser (633nm) for relevant fluorophore acquisition.

Identification of relevant structures by image analysis was determined using CellProfiler software (v2.8.0, The Broad Institute) ^52^. In the case of IMLS cell analysis for mitophagy, red only structures were identified by dividing the red signal by green signal per pixel following background noise reduction and weighting of the red signal to match that of the green signal in non-mitophagy inducing controls. By this method a value of ~1 indicates “yellow” networked mitochondria and values <1 represent mitochondria that have a stronger red signal than green signal. Values of <0.5 were taken to represent true red structures, regions that are therefore twice as bright for red than green.

### Zebrafish Maintenance and *in situ* hybridisation (ISH)

Wild-type zebrafish (AB strain) and transgenic tandem-tagged mitofish (TT-mitofish)^11^ were housed at the zebrafish facility at the Centre for Molecular Medicine Norway (AVD.172) using standard practices. Embryos were incubated in egg water (0.06 g/L salt (Red Sea)) or E3 medium (5 mM NaCl, 0.17 mM KCl, 0.33 mM CaCl2, 0.33 mM MgSO4, equilibrated to pH 7.0). Embryos were held at 28 °C in an incubator following collection. Experimental procedures followed the recommendations of the Norwegian Regulation on Animal Experimentation (“Forskrift om forsøk med dyr” from 15.jan.1996). All experiments conducted on wild-type zebrafish and transgenic tandem-tagged mitofish larvae were done at 5 dpf or earlier.

Whole-mount ISH for *prkcda* and *prkcdb* were performed as previously described using digoxigenin-labelled riboprobes^53^. Primer sequences for sense and antisense probes are described in Oligonucleotide Primers/Probes Appendix 2.

### Screening Library

Human targets for the siRNA library were identified by using the ExPASY PROSITE sequence motif database identifier for human proteins containing true-positive identified lipid binding domains. These included C1 domains (ID: PS50081), C2 domains (ID: PS50004), ENTH (ID: PS50942), PH Domain (ID: PS50003), PX Domain (ID: PS50195), FYVE domain (PS50178), GRAM or PROPPIN (SVP1 family) domains (No ID). This list was cross-checked against several previously published U2OS cell line proteomes (determined by mass spectrometry) and proteins not observed to be expressed in U2OS cells were removed ^15,16^. See Appendix 1 for a full list of siRNA targets.

### siRNA knockdown

The primary screen was carried out using a pooled siRNA approach with three Silencer Select siRNA oligonucleotides targeting each gene at 2.5nM final concentration each (7.5nM final). For transfection, 125μl of OptiMEM (Thermofisher Scientific #31985070) containing 100 ng/ml Doxycycline (Clontech #631311) and 0.1 μl RNAiMAX per pmol siRNA (Thermofisher Scientific #13778150) was added to each well of an Ibidi 96-well μ-plate (Ibidi #89626). After 5 mins at room temperature (RT), 25 μl of 75 nM siRNA (pooled) diluted in OptiMEM was added per well and incubated a further 15 mins at RT. U2OS IMLS cells were trypsinised and resuspended in complete media before centrifugation at 300 × g for 5 mins at RT. Media was removed and cells resuspended in OptiMEM to 2×10^5^ cells/ml. 100 μl of cells were added per well and samples were incubated for 16 h at 37°C. The media was then removed and changed to complete media for a further 24 h before the media again was changed to complete media ± 1mM DFP and incubated for 24h to induce mitophagy. Control wells with BafA1 treatment were dosed 2h prior to fixation. At the end of the experiment, samples were washed once in PBS and then fixed in 3.7% PFA, 200 mM HEPES pH 7 for 15mins /37°C. PFA was then quenched by washing twice and incubating a further 15mins in DMEM + 10mM HEPES pH 7. Wells were then washed twice with PBS and then incubated in PBS + 2 μg/ml Hoechst to stain nuclei for a minimum of 1h prior to imaging. Images were obtained on a Zeiss AxioObserver widefield microscope with a 20x objective acquiring a minimum of 35 fields of view per treatment. Analysis of red only punctate structures was carried out utilizing CellProfiler from a minimum of 1000 cells per condition per replicate.

### Identification and plotting of protein-protein interactome (PPI) networks

PPI represents the physical interaction among a set of proteins. PPI was obtained from Biological General Repository for interaction Datasets (BioGRID) version BIOGRID-ORGANISM-3.5.185.mitab^54^ (compiled April 25^th^ 2020) containing non-redundant and curated interactions. The networks were visualized using Cytoscape (v3.8.0) ^55^, we considered only the connected component of these seed networks for statistical and functional analysis. Functional and pathway analysis of connected components of interaction network was performed by ShinyGO^56^. We only considered GO terms for cellular component, molecular functions and biological process with significant p-value and enrichment values. Graphs were plotted using R package ggplots.

### RNA Isolation, cDNA synthesis and qPCR

For quantifying knockdown in the secondary deconvolution siRNA screen, RNA was isolated and cDNA generated from transfected U2OS cells using *Power* SYBR Green Cells-to-C_T_ kit (ThermoFisher Scientific #4402955) as per manufacturer’s instructions.

For other experiments, RNA was isolated from cells or zebrafish (~50 embryos per sample) using Trizol reagent (ThermoFisher Scientific #15596026). cDNA was synthesised from RNA using Superscript III reverse transcriptase (ThermoFisher Scientific #18080085) according to manufacturer’s instructions. Amplification was performed with KAPA SYBR FAST qPCR Kit using a CFx96 real-time PCR system (Bio-Rad) using primers designed to amplify target genes as indicated in Oligonucleotide Primers/Probes

Appendix 2 following normalisation of transcript levels to TATA-box-binding protein (TBP – cell samples) or β-actin (zebrafish samples) using the 2^−ΔΔCt^ method.

### Western Blotting

For western blotting experiments, cells were treated as indicated in figure legends prior to moving onto ice and washing twice with cold PBS. Cells were lysed on ice in NP-40 lysis buffer [50mM HEPES pH 7.4, 150mM NaCl, 1mM EDTA, 10% (v/v) Glycerol, 0.5% (v/v) NP-40 + 1mM DTT, 1x Phosphatase inhibitors and 1x Protease inhibitors fresh] and incubated 5 mins prior to collecting. For zebrafish samples, embryos were collected at 3 dpf and lysed in RIPA buffer [50 mM Tris-HCl pH 8, 150 mM NaCl, 5 mM EDTA, 1 % NP-40, 0.5 % Sodium deoxycholate, 0.1 % SDS, 1x protease inhibitor cocktail], approximately 20-30 embryos were used per gel lane.

Samples were clarified by centrifugation at 21000 × g / 4 °C / 10 mins and supernatant retained. Protein levels were quantified by Bradford assay (Bio-Rad #5000006) relative to a BSA standard. Samples were normalised and added to loading sample [1x = 62.5mM Tris pH 6.8, 10% (v/v) Glycerol, 2% (w/v) SDS, 0.005% (w/v) Bromophenol Blue] to achieve 30μg-50μg of protein per lane. Samples were ran by acrylamide gel and transferred to PVDF (350mA/50mins). Samples were blocked in TBS Odyssey Blocking Buffer (Li-Cor #927-50000) for 30mins/RT before incubation overnight at 4°C with primary antibodies (TBS blocking buffer + 0.2% Tween). Membranes were washed 3×10 mins in TBST before secondary antibody incubation (TBS blocking buffer + 0.2% Tween + 0.01% SDS) for 1h. Membranes were washed 3x 10min with TBST before a final wash in TBS only and membrane imaging.

### Antibodies

Primary antibodies targeting ATG13 (#13468, Clone E1Y9V), AMPK P-T172 (#2535), β-Actin (#3700, Clone 8H10D10), BNIP3 (#44060, Clone D7U1T), BNIP3L (#12396, Clone D4R4B), COXIV (#4850, Clone 3E11), PDH (#2784), PRKCD (#9616, D10E2), PRKCD P-S663 (#9376), LC3B (Western blotting only, #3868, Clone D11), ULK1 (#8054), ULK1 P-S555 (#5869), ULK1 P-S757 (#6888, Clone D1H4) were from Cell Signaling Technology. FUNDC1 (#Ab74834), GAK (#Ab115179, Clone 1C2), NIPSNAP1 (#Ab67302), MTCO2 (#Ab110258, Clone 12C4F12) and WIPI2 (#Ab105459, Clone 2A2) were from Abcam. ATG13 P-S318 (#600-401-C49S) was from Rockland. HIF1α (MAB1536-SP, Clone 241809) was from R&D Systems. LAMP1 (sc-20011, Clone H4A3) was from Santa Cruz Biotechnology. α-Tubulin (T5168, Clone B-5-1-2) was from Sigma Aldrich.

Secondary antibodies for western blotting are indicated in the source data file, these included anti-rabbit (Starbright Blue, Bio-Rad, 12004161) (DyLight 800, ThermoFisher Scientific, SA5-10044) (DyLight 680, ThermoFisher Scientific, SA5-10042) or anti-mouse (Starbright Blue, Bio-Rad,12004158) (DyLight 680, ThermoFisher Scientific, SA5-10170) or anti-tubulin (hFAB Rhodamine, Bio Rad, 12004166). Secondary antibodies for immunofluorescence were anti-rabbit Alexa Fluor-594 (Invitrogen, A11058), Alexa Fluor-647 (ThermoFIsher Scientific, A21245) or anti-mouse Alexa Fluor-647 (ThermoFisher Scientific, A21236).

### Phos-Tag Gels

Phos-tag acrylamide gels were prepared in line with manufacturer’s instructions. Briefly, 8% resolving poly-acrylamide gels were prepared containing 25μM Phos-tag reagent (Wako Chemicals #AAL-107) and 50μM MnCl_2_ 26. Samples to be ran for analysis were diluted in loading sample containing 10 mM MnCl_2_. Acrylamide gels were ran at 40 mA until complete and washed 3×10 min/RT in 1x transfer buffer (48mM Tris, 39mM Glycine, 0.0375% (w/v) SDS) + 10mM EDTA followed by 1×10min in 1x transfer buffer. Samples were then transferred to PVDF at 350mA / 50min and treated as noted earlier for western blot samples.

### PFA Fixation, antibody staining and imaging

Cells to be imaged were seeded onto glass coverslips 16h prior to treatments as indicated in figure legends. Following treatment, cells were washed once with PBS prior to addition of warmed fixation buffer (3.7% (w/v) PFA, 200mM HEPES pH 7.4) or for double tag IMLS cells (3.7% (w/v) PFA, 200mM HEPES pH 7) and incubated 15 mins at 37°C. Coverslips were washed twice and incubated 1×15 mins with DMEM + 10mM HEPES pH7.4 (IMLS = pH 7). Cells were then washed once with PBS and then permeabilised by incubation for 5 mins with permeabilisation buffer (0.2% (v/v) NP-40 in PBS). Cells were washed twice and then incubated 20 mins with IF blocking buffer (PBS + 1% (w/v) BSA) to block the samples. Coverslips were then incubated 1 h / 37 °C with primary antibodies diluted in IF blocking buffer before washing 3×10 mins in IF blocking buffer. Coverslips were then incubated 30 mins / RT with appropriate secondary antibodies. Finally, samples were washed 3×10 mins in IF blocking buffer prior to mounting on to coverslides with ProLong Diamond Antifade Mountant with DAPI (ThermoFisher Scientific #P36962). Slides were allowed to cure overnight before imaging with either a Zeiss AxioObserver widefield microscope (20x) or Zeiss LSM 800 confocal microscope (60x).

### Citrate Synthase Assay

To biochemically quantify mitochondrial abundance, we assayed citrate synthase activity from cell lysates. Briefly, U2OS cells were grown and subject to treatments as described in figure legends, cells were then washed twice with PBS on ice before lysis [50 mM HEPES pH 7.4, 150 mM NaCl, 1 mM EDTA, 10 % Glycerol, 0.5 % NP-40, 1 mM DTT, 1x Phosphatase inhibitors, 1x Protease inhibitors]. Cell lysates were clarified by centrifugation at 21000 × g / 10 min / 4 °C and supernatants retained. Protein concentration was determined by Bradford assay. To determine citrate synthase activity 1 μl of protein lysate was added to 197 μl of CS assay buffer [100 mM Tris pH 8, 0.1 % Triton X-100, 0.1 mM Acetyl CoA, 0.2 mM DTNB [5,5’Dithiobis(2-nitrobenzoic acid)]) in a multi well plate. At the assay start point, 2 μl of 20 mM Iodoacetamide was added per well and incubated at 32 °C and reactions monitored at λ_Abs_=420 nm for 30 min in a FLUOstar OPTIMA (v2.20R2, BMG Labtech) plate reader and compared to iodoacetamide null controls. The Δ λ_Abs_ was plotted and the reaction rate determined across the linear range before saturation. The reaction rate was then normalised to the protein concentration and plotted relative to the control.

### Correlative Light Electron Microscopy (CLEM)

For CLEM, U2OS IMLS cells were grown on photo-etched coverslips (Electron Microscopy Sciences, Hatfield, USA). The next day, cells were treated with DFP (1mM) ± GAKi (10 μM) for 24 h. Cells were then fixed in 4 % formaldehyde, 0.1 % glutaraldehyde/0.1 M PHEM (60 mM PIPES, 25 mM HEPES, 2 mM MgCl_2_, 10 mM EGTA, pH 6.9), for 1 h. The cells were mounted with Mowiol containing 2 μg/ml Hoechst 33342 (Sigma-Aldrich). Mounted coverslips were examined with a Zeiss LSM710 confocal microscope with a Zeiss plan-Apochromat 63x/1.4 Oil DIC III objective. Cells of interest were identified by fluorescence microscopy and a Z-stack was acquired. The relative positioning of the cells on the photo-etched coverslips was determined by taking a DIC image. The coverslips were removed from the object glass, washed with 0.1 M PHEM buffer and fixed in 2 % glutaraldehyde/0.1 M PHEM for 1h. Cells were post fixed in osmium tetroxide and uranyl acetate, stained with tannic acid, dehydrated stepwise to 100% ethanol and flat-embedded in Epon. Serial sections (~100-200nm) were cut on an Ultracut UCT ultramicrotome (Leica, Germany), collected on formvar coated slot-grids. Samples were observed in a Thermo ScientificTM TalosTM F200C microscope and images were recorded with a Ceta 16M camera. For tomograms, single-tilt image series were recorded between −60° and 60° tilt angle with 2° increment. Single axis tomograms were computed using weighted back projection and, using the IMOD software package version 4.9^57^.

### Mitochondrial Enrichment

Cells to be enriched for mitochondria were grown and treated as noted in figure legends. Cells were then moved to ice and washed twice with ice cold PBS. 1 ml of mito fractionation buffer (5 mM Tris-HCl pH 7.5, 210 mM Mannitol, 70 mM Sucrose, 1 mM EDTA pH 7.5, 1 mM DTT, 1x protease and phosphatase inhibitors) was added per 10 cm dish and scraped to collect cells. A “cell homogenizer” (Isobiotec) was utilised with a 16 μm clearance ball and prepared by passing through 1 ml of mito fractionation buffer. Cell solution was collected in a 1 ml syringe and passed through the cell homogenizer 9 times. The resulting mix was centrifuged 500 × g / 4°C / 5 mins to pellet unbroken cells and nuclei. The supernatant was taken, and a small sample retained as post nuclear supernatant, the remaining was centrifuged at 10000 × g/4°C/ 10 mins to pellet mitochondria. The supernatant was removed to waste, and the pellet resuspended in 500 μl mito fractionation buffer and 10000 × g/4 °C/10 mins centrifugation step repeated. The supernatant was removed once more, the pellet represents enriched mitochondria that could be added directly to protein loading sample for downstream western blotting.

### Mitochondrial Classifier

Cellular mitochondria were classified as tubular or fragmented by implementing an image classified utilising CellProfiler Analyst (v2.2.1, The Broad Institute). Classifications were determined by using siDRP1 and siOPA1 treated cells as positive controls for fragmented and hyperfused phenotypes respectively. Classifier was trained on the EGFP fluorescent images and with a confusion matrix of >0.90 for each phenotype.

### Crystal Violet Staining

U2OS cells were seeded into 96-well plates in triplicate at 2×10^4^ cells per well and incubated overnight in complete media. Cells were then treated for 24 h with indicated compounds and doses, utilising puromycin as a positive control. Following treatment, cell media was removed and cells washed twice with a gentle stream of water. This was then removed and 100 μl of staining solution (0.5% (w/v) crystal violet, 20% methanol) added and incubated 20 min / RT with gentle rocking. Wells were washed 4x with water, all liquid removed and left overnight to air dry. Then 200μl per well of 100% methanol for 20min/RT was added with gentle rocking and sample absorbance read at OD_570_. Sample values were adjusted by no-well control (blank) wells and viability determined by normalisation to an untreated control.

### Quantification of mitophagy in *C. elegans*

The strain used to monitor mitophagy process in *C. elegans* was IR2539:*unc-119(ed3);Ex*[_pmyo-_3TOMM-20::Rosella;unc-119(+)]. Standard procedures for *C. elegans* strain maintenance were followed. Nematode rearing temperature was kept at 20°C. For RNAi experiments worms were placed on NGM plates containing 2 mM IPTG and seeded with HT115(DE3) bacteria transformed with either the pL4440 vector or the gakh-1 RNA construct for two generations. Synchronous animal populations were generated by hypochlorite treatment of gravid adults to obtain tightly synchronized embryos that were allowed to develop into adulthood under appropriate, defined conditions. Progeny of these adults were tested on adult day 2. We performed imaging of mitophagy process in *C. elegans* based on the methods we had established ^58–60^. Briefly, worms were immobilized with levamisole before mounting on 2% agarose pads for microscopic examination using EVOS Imaging System. Images were acquired as Z-stacks under the same exposure. Average pixel intensity values and frequency of GFP/DsRed puncta were calculated by sampling images of different animals. The calculated mean pixel intensity for each animal in these images was obtained using FIJI.

### CRISPR/Cas9 genome editing in zebrafish and microinjections

To generate *prkcda* and *prkcdb* knock-out embryos, we utilised CRISPR/Cas9 as described earlier (Jao et al., 2013). Potential gRNA target sites were identified using the online web tool CRISPR Design (http://CRISPR.mit.edu) or CHOPCHOP (http://chopchop.cbu.uib.no/index.php) (Montague et al., 2014). Genomic DNA sequences retrieved from Ensembl GRCz10 or z11 (http://uswest.ensembl.org/Danio_rerio/Info/Index) were used for the target site searches. Three guide RNAs were designed each for *prkcda* and *prkcdb* respectively, based on predictions from the aforementioned web programs. All sgRNAs were prepared by *in vitro* transcription of double-stranded deoxyoligonucleotide templates as described previously ^61^. Cas9 nuclease (EnGen Cas9 NLS, NEB) was combined with an equimolar mixture of 3x sgRNA’s (or 6x for *prkcd_ab* DKO) and incubated for 5-6 minutes at room temperature. After incubation, the mixture was immediately placed back on ice, until pipetted into the capillary needle used for microinjection and then approximately 1 nl of 5 μM sgRNA:Cas9 complex was microinjected into the cytoplasm of one celled stage zebrafish embryo.

Oligonucleotides used for sgRNA synthesis are listed in Oligonucleotide Primers/Probes Appendix 2, a universal primer was used with individual sgRNA primers (5’AAAAGCACCGACTCGGTGCCACTTTTTCAAGTTGATAACGGACTAGCCTTA TTTTAACTTGCTATTTCTAGCTCTAAAAC’3).

### Zebrafish Locomotor Assay

Larval motility was monitored using the ZebraBox and Viewpoint software (v3.10.0.42, Viewpoint Life Sciences Inc) under infrared light. At 5 days post fertilization (dpf), larvae were singly placed in 48-well plates with 300 μl of fish water per well, followed by incubation at 28.5 ^o^C on a normal light cycle overnight. All experiments were completed in a quiet room at 5 dpf between 10 AM and 2 PM. Larvae were allowed to acclimate in the ZebraBox measurement apparatus for 2 h before recording. Larvae were then exposed to alternating cycles of infrared light and dark, every 30 min as described ^63^. Larval locomotion was tracked with the Viewpoint software. Motility was defined as tracks moving less than 10 cm/s, but more than 0.1 cm/s.

### Zebrafish Mitophagy cryosectioning and confocal microscopy

Zebrafish mitophagy experiments were conducted on tt-mitofish with relevant prkcd_a/b KO lines as described above. To examine mitophagy, zebrafish larvae were treated with DMOG or control for 24 h. At experimental end-points, larvae were washed once in embryo water and fixed with 3.7% PFA (in HEPES, pH 7-7.2) at 4 °C overnight. Post fixation, larvae were washed three times in PBS. The larvae were then cryopreserved in a 2 mL tube in increasing amounts of sucrose in 0.1 M PBS with 0.01 % sodium azide. Cryopreservation was done first in 15 % sucrose solution for 1 hour at RT or up until the larvae drops to the bottom of the tube and then in 30% sucrose solution at 4 °C overnight with gentle shaking. Cryopreserved larvae were oriented in a cryomold (Tissue-Tek Cryomold, Sakura, Ref: 4565) with optimal cutting temperature compound (OCT compound) (Tissue-Tek Sakura, Ref: 4583). Larvae were oriented with the ventral side down and additional OCT was added to fill the mold and frozen down on dry ice. A solid block of OCT with couple of larvae oriented in the desired way, was taken out from the mold and 12 μm coronal slices were sectioned on the cryostat (Thermo Scientific). Sections were collected on Superfrost Plus glass slides (Thermo Scientific, Ref: J1800AMNZ) and kept at RT for at least 2 h to firmly tether slices onto the glass slide.

The pH of all solutions and buffers used were 7-7.2. Slides were rehydrated three times in PBST (0.1 % Tween 20 in 1X PBS) at room temperature for 3 minutes each. Area of interest was circled by a hydrophobic PAP pen (Abcam, ab2601) and the slides were placed in a humidified chamber to avoid drying out. 100-200 μl of 1 μg/ml Hoechst solution was gently pipetted onto the slides and incubated for 30 mins at RT. Post incubation, slides were washed 3 times in PBST for 5 minutes each and mounted using ProLong Diamond Antifade Mountant (Invitrogen, P3696). Coverslips were carefully placed over the sections. Confocal images were obtained using an Apochromat 20x/0.8 or 63x/1.2 oil DIC objective on an LSM 800 microscope (Zeiss). Red puncta were counted manually from hind-brain regions.

### Mass Spectrometry

#### Sample Preparation

Cells were dissolved in RIPA buffer and further homogenized with a sonicator (30 sec × 3 times with 30 sec interval) and insoluble material was removed by centrifugation. Protein concentrations were estimated by BCA assay (Pierce). For each replicate, 600 μg of protein samples for phosphoproteomics and 30 μg for whole cell lysate proteomics were reduced and alkylated and further digested with trypsin by FASP (Filter aided sample preparation) method. Digested peptides were transferred to a new tube, acidified and the peptides were de-salted using Oasis cartridges for STY peptides enrichments. Phosphorylated peptides enrichment was performed based on TiO_2_ 64. Enriched peptides fractions were de-salted by C_18_ stage tips.

#### LC-MS/MS

Peptide samples were dissolved in 10 μl 0.1 % formic buffer and 3 μl were loaded for MS analysis. The Ultimate 3000 nano-UHPLC system (Dionex, Sunnyvale, CA, USA) connected to a Q Exactive mass spectrometer (ThermoElectron, Bremen, Germany) equipped with a nano electrospray ion source was used for analysis. For liquid chromatography separation, an Acclaim PepMap 100 column (C18, 3 μm beads, 100 Å, 75 μm inner diameter) (Dionex, Sunnyvale CA, USA) capillary of 50 cm bed length was used. A flow rate of 300 nL/min was employed with a solvent gradient of 3-35 % B in 220 mins, to 50 % B in 20 min and then to 80 % B in 2 min. Solvent A was 0.1 % formic acid and solvent B was 0.1 % formic acid/90% acetonitrile.

The mass spectrometer was operated in the data-dependent mode to automatically switch between MS and MS/MS acquisition. Survey full scan MS spectra (from m/z 400 to 2000) were acquired with the resolution R = 70,000 at m/z 200, after accumulation to a target of 1e6. The maximum allowed ion accumulation times were 100 ms. The method used allowed sequential isolation of up to the ten most intense ions, depending on signal intensity (intensity threshold 1.7e4), for fragmentation using higher collision induced dissociation (HCD) at a target value of 10,000 charges and a resolution R = 17,500. Target ions already selected for MS/MS were dynamically excluded for 30 sec. The isolation window was m/z = 2 without offset. The maximum allowed ion accumulation for the MS/MS spectrum was 60 ms. For accurate mass measurements, the lock mass option was enabled in MS mode and the polydimethylcyclosiloxane ions generated in the electrospray process from ambient air were used for internal recalibration during the analysis.

#### Data Analysis

Raw files from the LC-MS/MS analyses were submitted to MaxQuant (v1.6.1.0) software for peptide/protein identification^65^. Parameters were set as follow: Carbamidomethyl (C) was set as a fixed modification; protein N-acetylation and methionine oxidation as variable modifications and PTY. A first search error window of 20 ppm and mains search error of 6 ppm was used. Minimal unique peptides were set to one, and FDR allowed was 0.01 (1 %) for peptide and protein identification. The Uniprot human database was used. Generation of reversed sequences was selected to assign FDR rates. MaxQuant output files (proteinGroups.txt for proteomic data and STY(sites).txt for phosphoproteomic data) were loaded into the Perseus software^66^. Identifications from potential contaminants and reversed sequences were removed and intensities were transformed to log2. Identified phosphorylation sites were filtered only for those that were confidently localized (class I, localization probability ≥ 0.75). Next, proteins identified in two out three replicates were considered for further analysis. All zero intensity values were replaced using noise values of the normal distribution of each sample. Protein or STY abundances were compared using LFQ intensity values and a two-sample Student’s T-test (permutation-based FDR correction (250 randomizations), FDR cut-off: 0.05, S0: 0.1).

The complete datasets have been uploaded to ProteomXchange.

### Statistics and Significance

Experimental values were used for statistical analysis using Prism (v8.0.1) where indicated using analyses and post-hoc tests as indicated in figure legends. All data values come from distinct samples. Where shown **** = p>0.0001, *** = p >0.001, ** = p>0.01, * = p > 005 or n.s = not significant.

## ACKNOWLEDGEMENTS

We would like to thank Coen Campersteijn for assistance with live cell imaging. We would also like to thank the Simonsen lab for their support and critical discussion throughout.

This work was supported by the Norwegian Cancer Society (Project: 171318) and the Research Council of Norway through its Centres of Excellence funding scheme (Project: 262652) and FRIPRO grant (Project: 221831).

## AUTHOR CONTRIBUTIONS

Experimental planning, data analysis and writing of the manuscript were performed by M.J.M and A.S with input from all authors. B.J.M carried out *D.rario* experiments, Seb.S carried out CLEM experiments. Y.A and E.F carried out *C.elegans* experiments. Sac.S and J.W prepared and ran samples for MS analysis. Sak.S carried out analysis of interaction networks. A.H.L generated the IMLS cell line. M.J.M, L.T.M, M.Y.W.N.G and L.R.dlB performed all remaining experiments.

## AUTHOR DECLARATIONS

M.J.M is now an employee of AstraZeneca plc.

## DATA AVAILABILITY

All data is available upon reasonable request. Source data for quantitative figures and supplementary figures is provided as supplementary information. Proteomics Data has been uploaded to PRIDE.

## APPENDICES

## siRNA Targets List

**Appendix 1.**
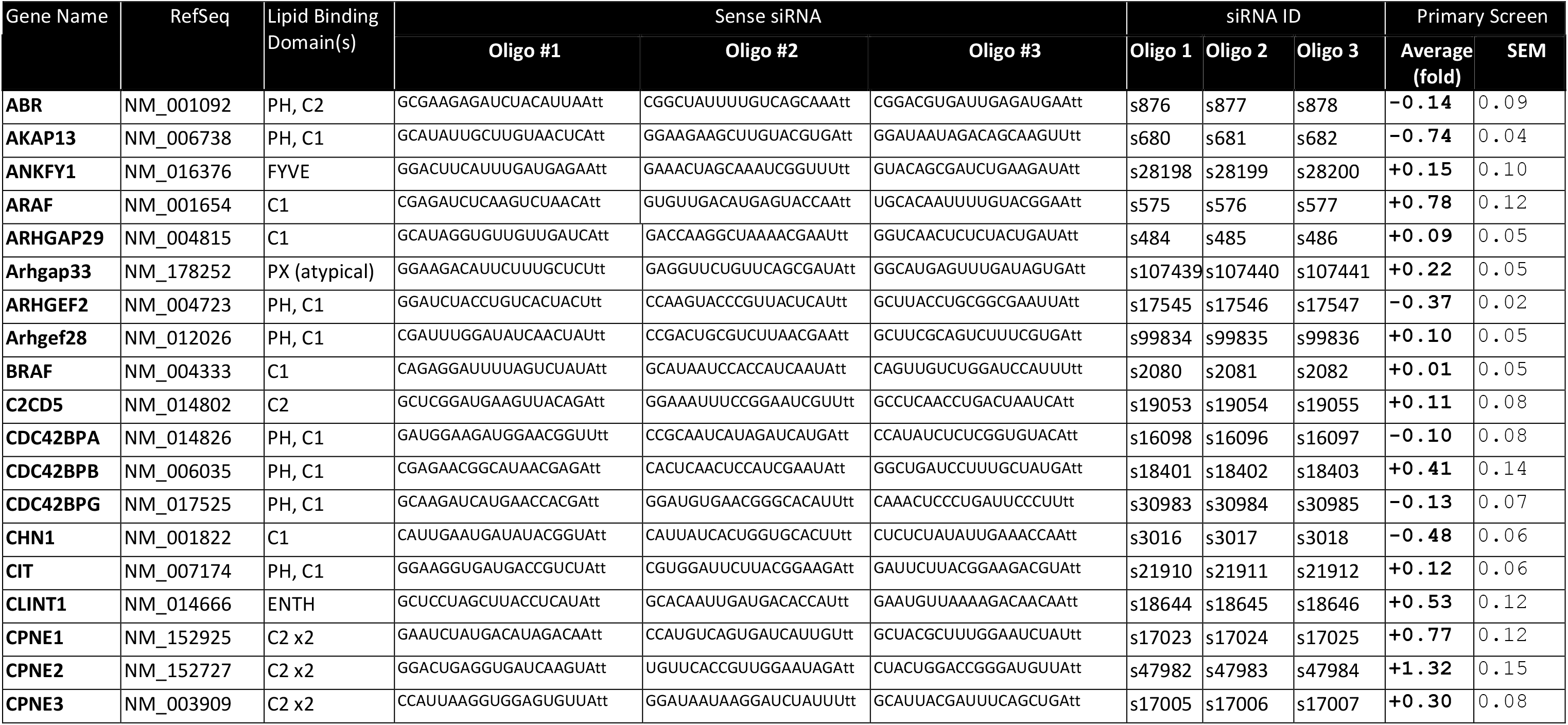
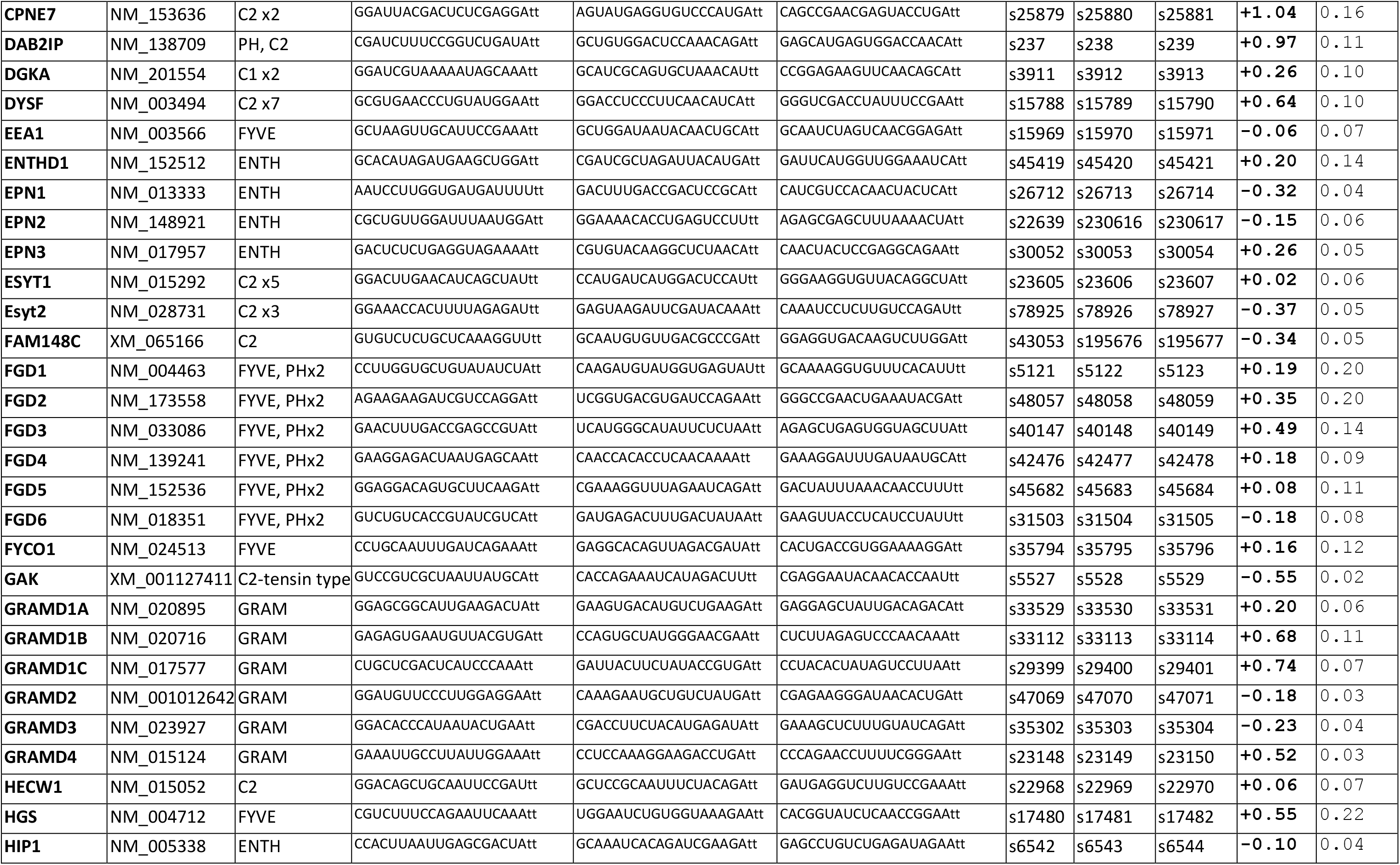
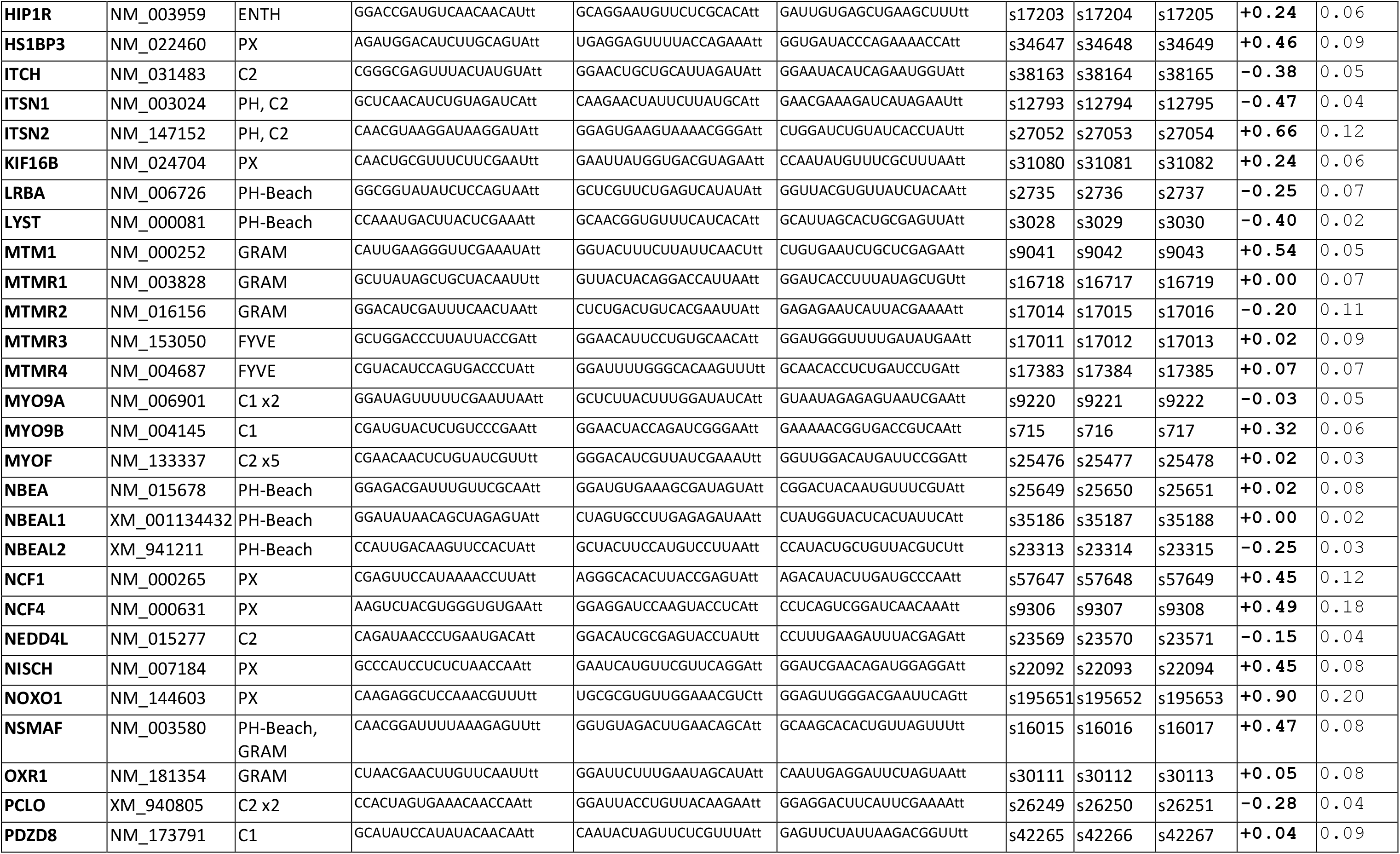
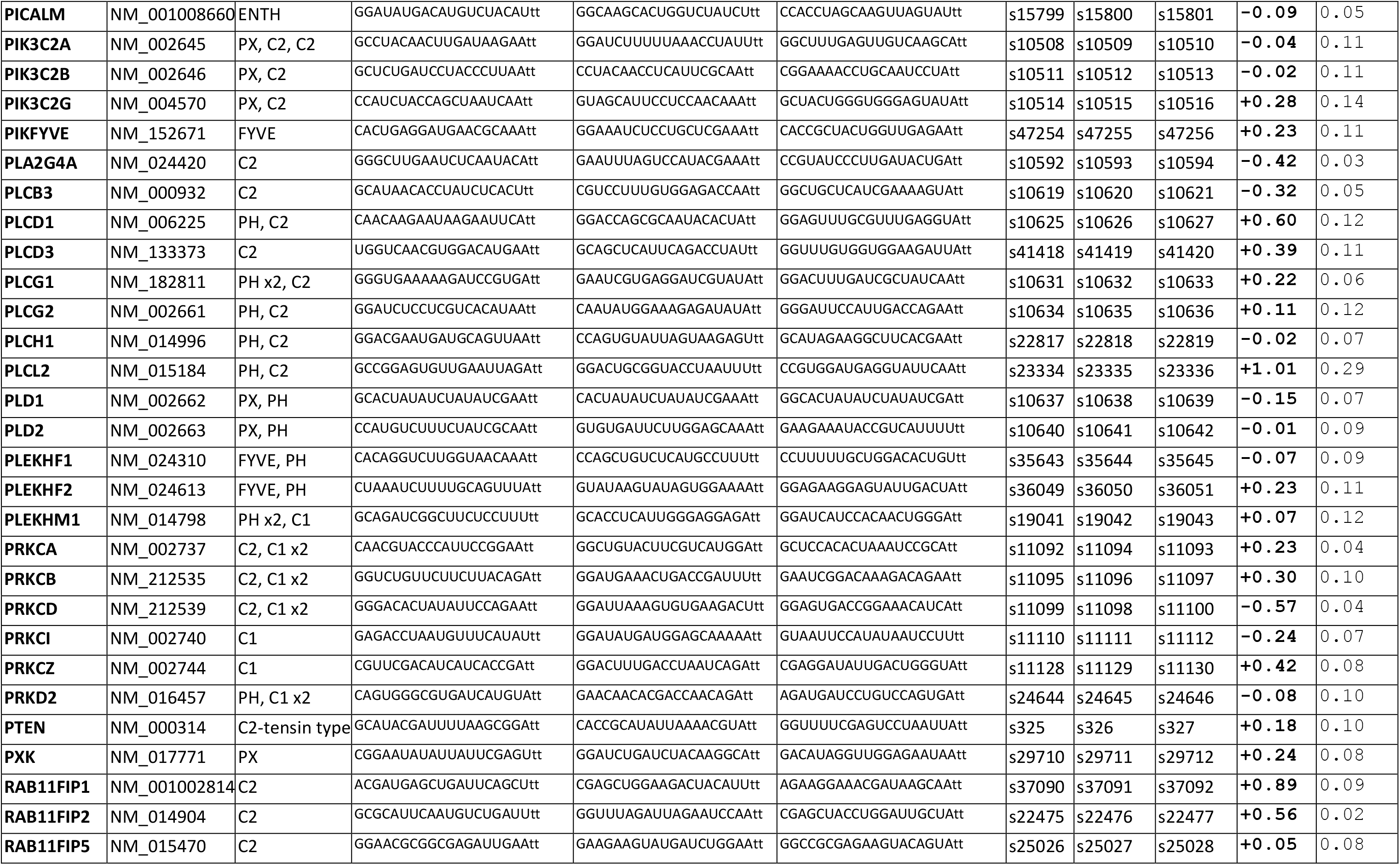
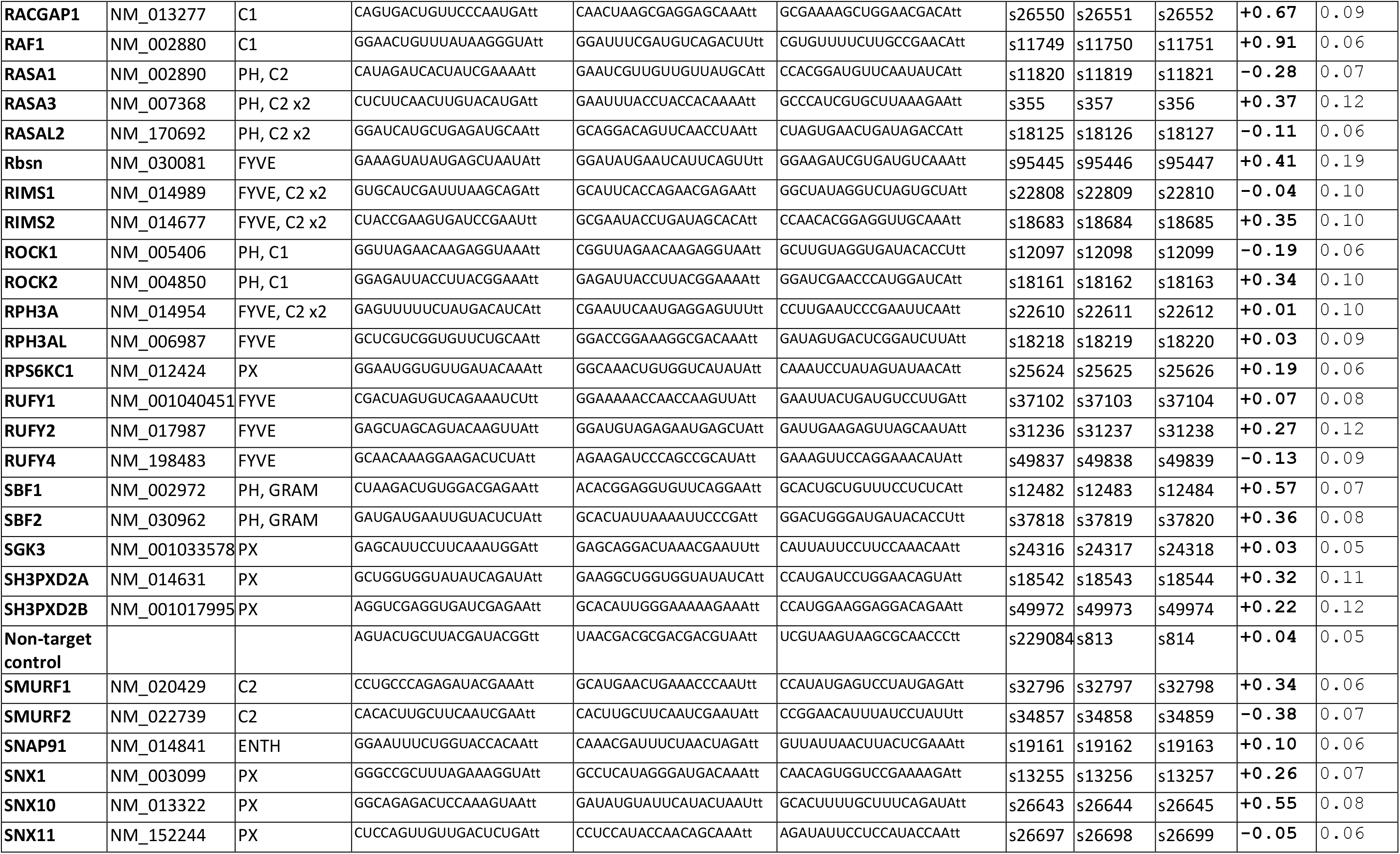
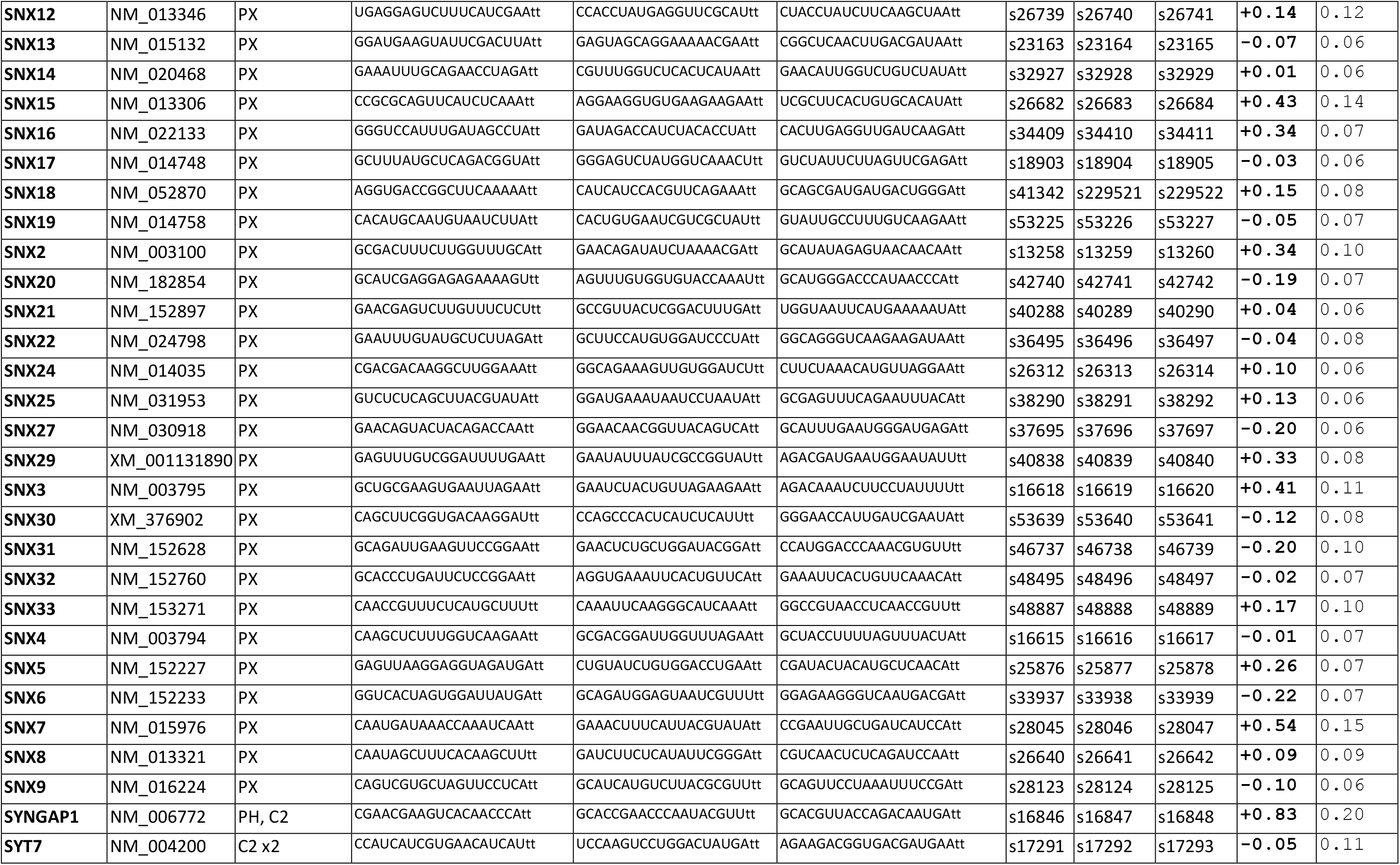
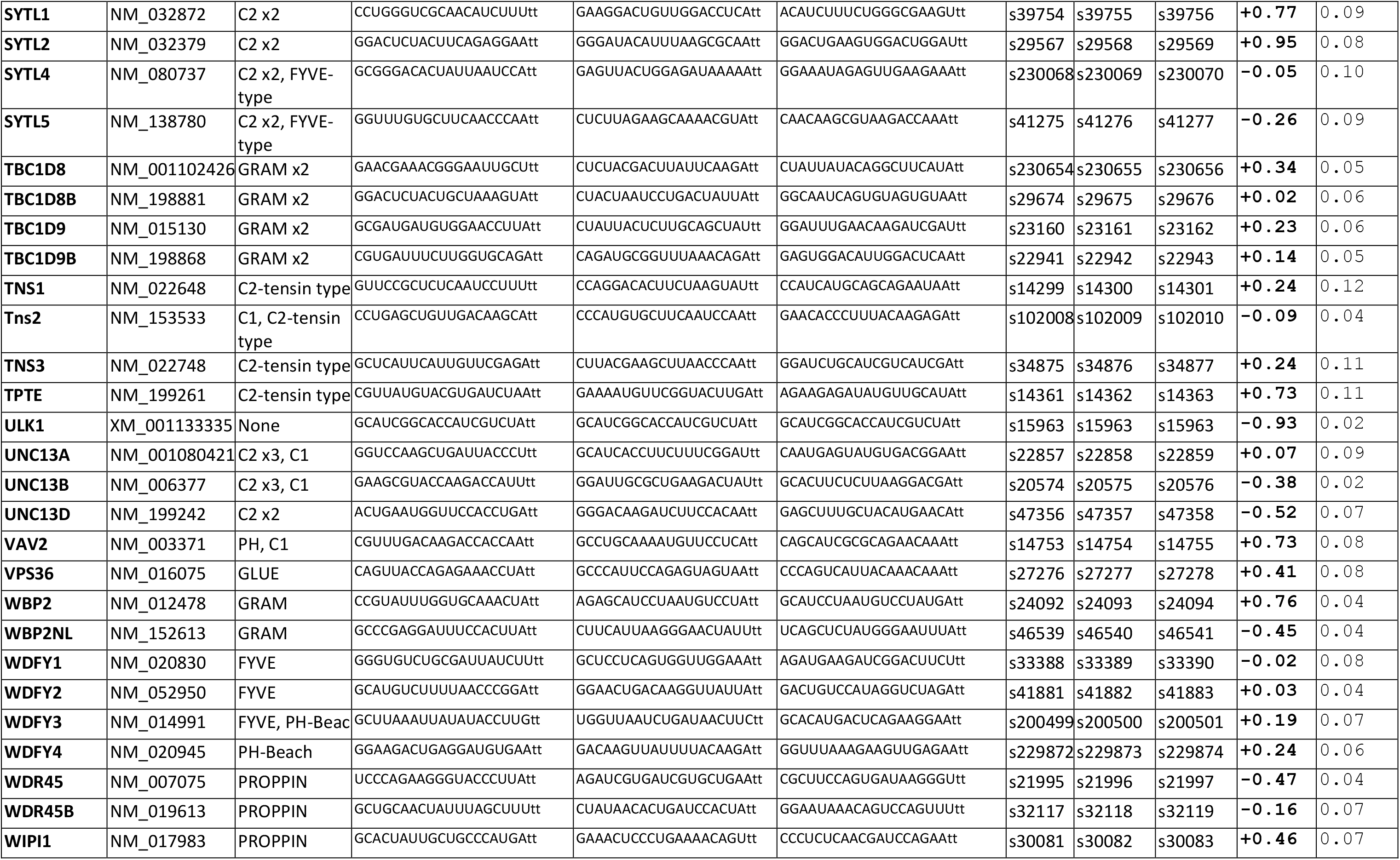
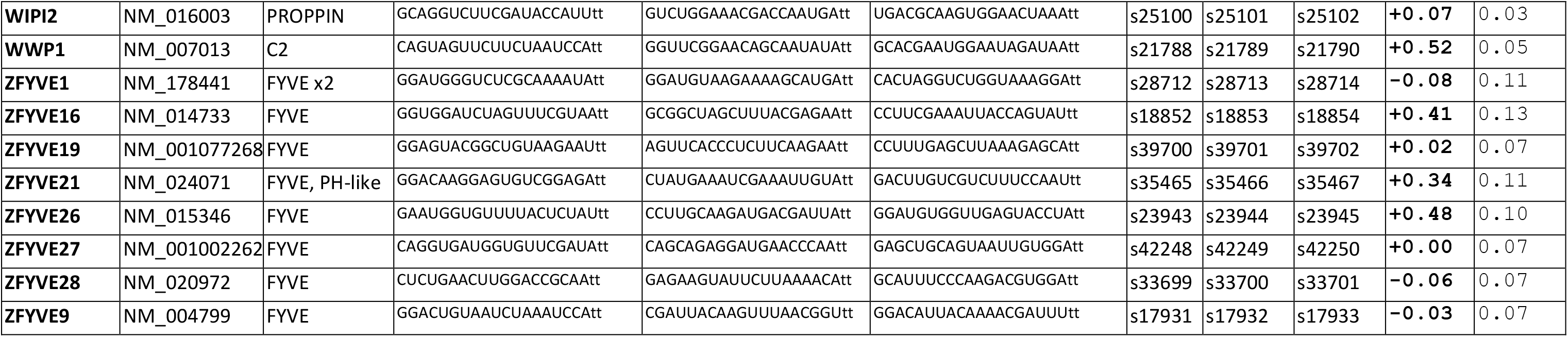
siRNA Targets.

## Oligonucleotide primers/probes

**Appendix 2.**
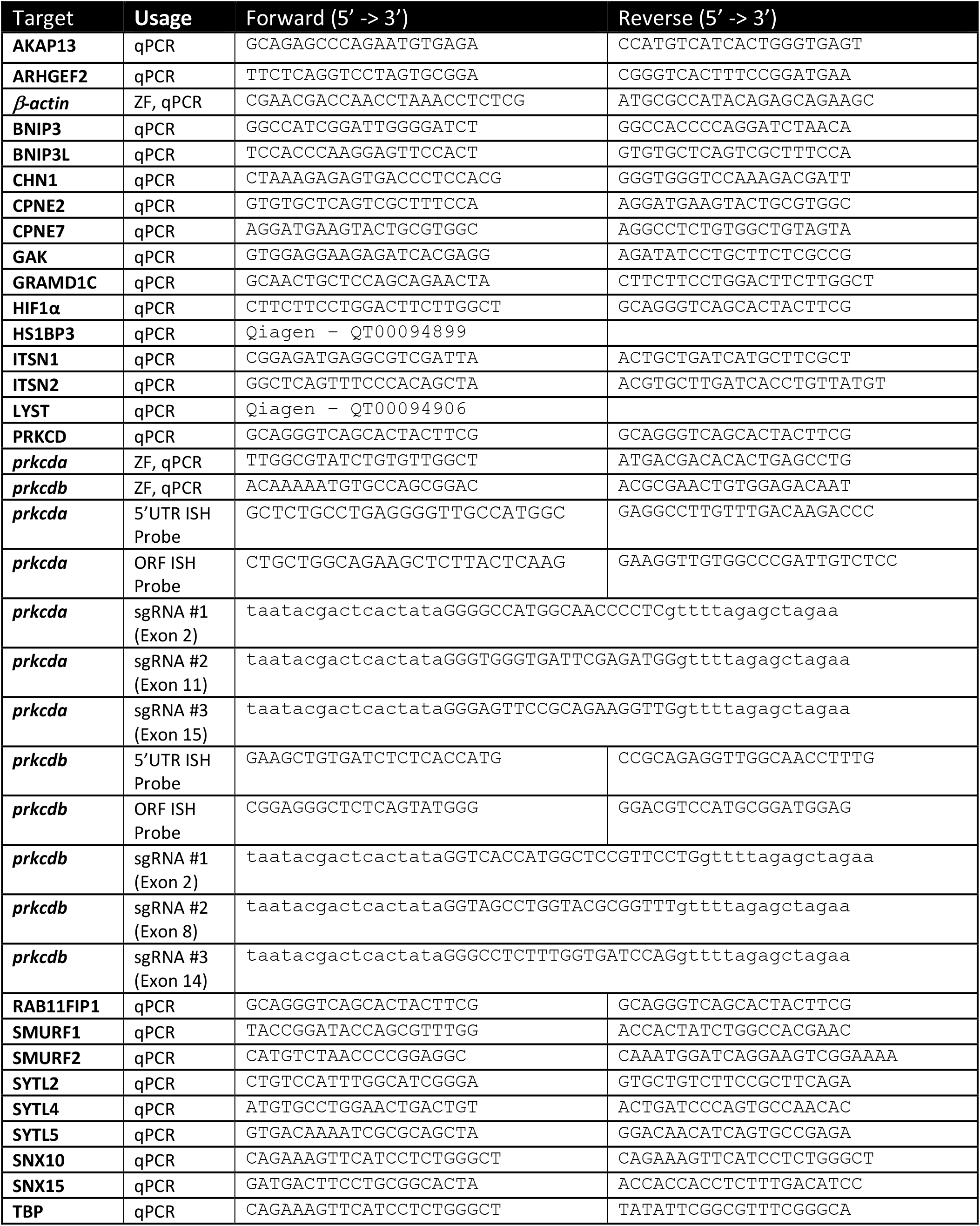
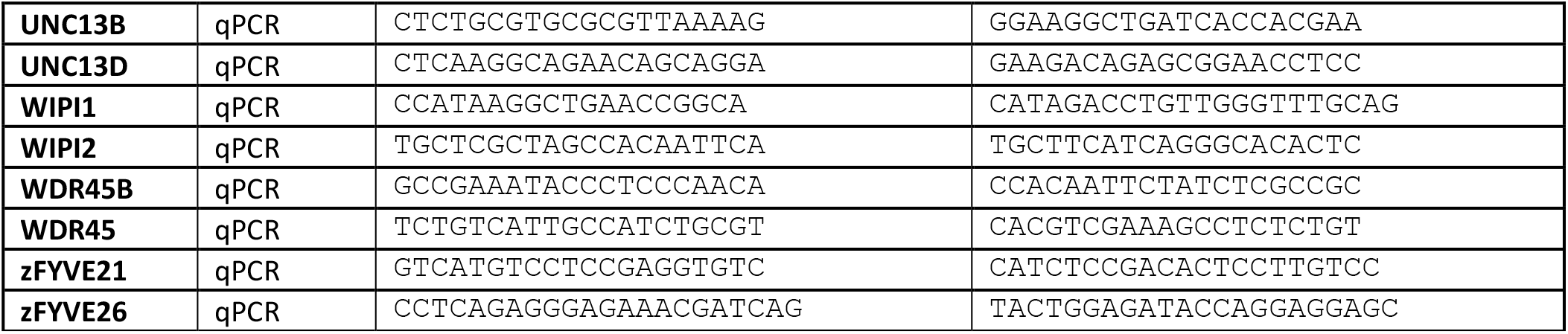
Oligonucleotide primers/probes.

