## Supplemental Figures for "GAK and PRKCD are positive regulators of PRKN-independent mitophagy"

### Supplementary Table 1

| Protein | Localisation | Change ($Log_2$) |
|---|---|---|
| TOMM70A | OMM | -0.11 |
| TOMM34 | OMM | -0.19 |
| VDAC2 | OMM | +0.13 |
| MRPL16 | Matrix | -0.62 |
| MRPL27 | Matrix | -1.76 |
| COQ9 | Matrix | -0.87 |
| TIMM23 | IMM | -1.51 |
| TIMM8B | IMM | -0.74 |
| NDUFA12 | IMM | -1.28 |

**Supplementary Table 1 - A selection of mitochondrial proteins and their intensity change (Log2) between control and DFP treatments from Fig. 1e**

### Supplementary Figure 1

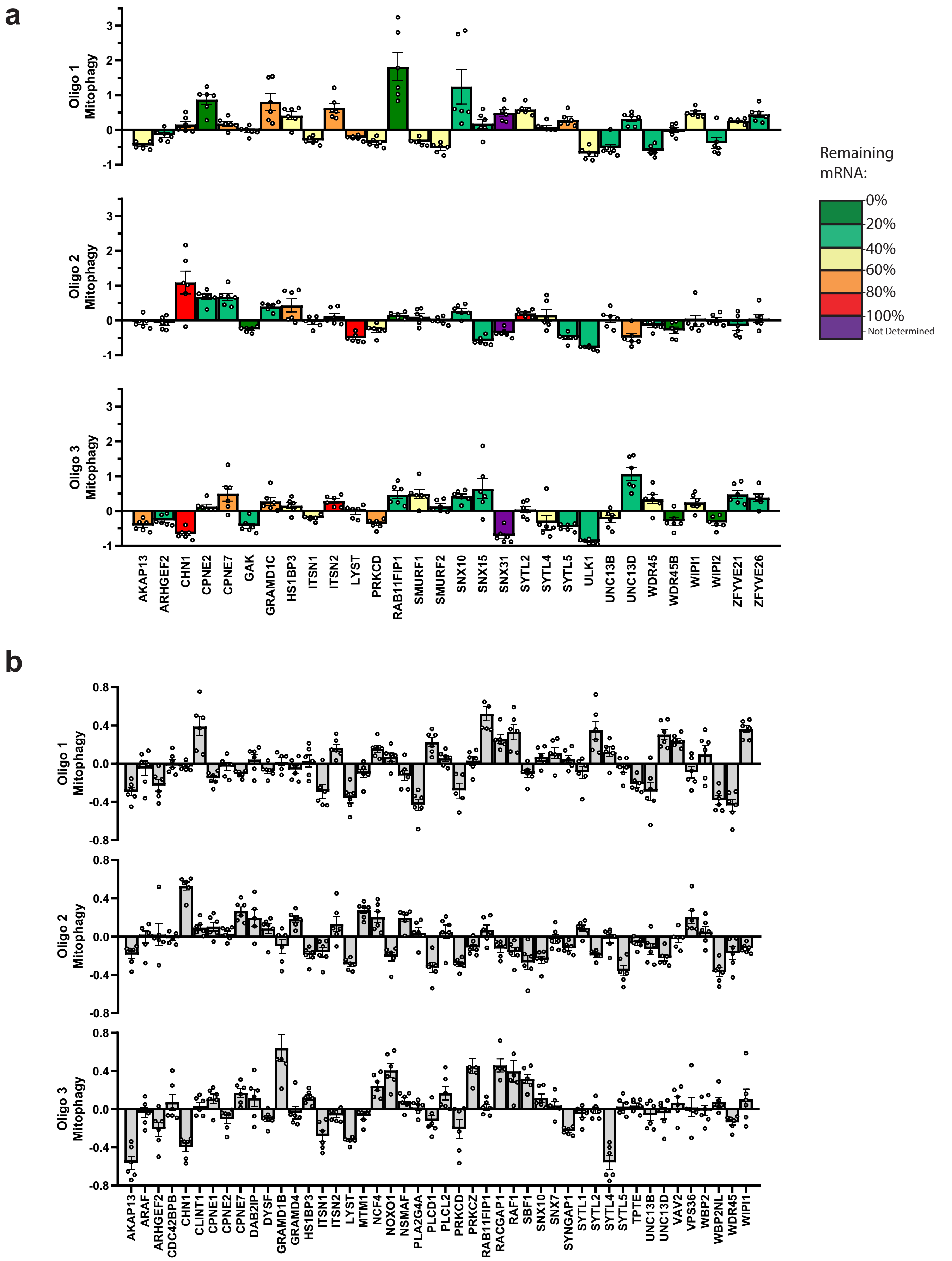


**Supplementary Fig. 1 – Secondary and Tertiary siRNA screen**

**a** Secondary screen of targets identified in the primary screen, carried out by transfection of the individual single siRNA oligos (7.5 nM each) for 48 h prior to 24 h 1 mM DFP treatment. Bars represent mean fold change in mitophagy relative to the siNT controls ± SEM from n=6 plates, colour represents the level of mRNA knockdown ascertained by qPCR analysis where green = high knockdown and red = poor knockdown. **b** Tertiary siRNA screen carried out by transfection of individual siRNA oligos (15 nM each) for 72 h prior to 24 h 1 mM DFP treatment. Bars represent mean fold change in mitophagy relative to the siNT controls ± SEM from n= 6 plates.

### Supplementary Figure 2

**a**

#### GAK Interaction Map

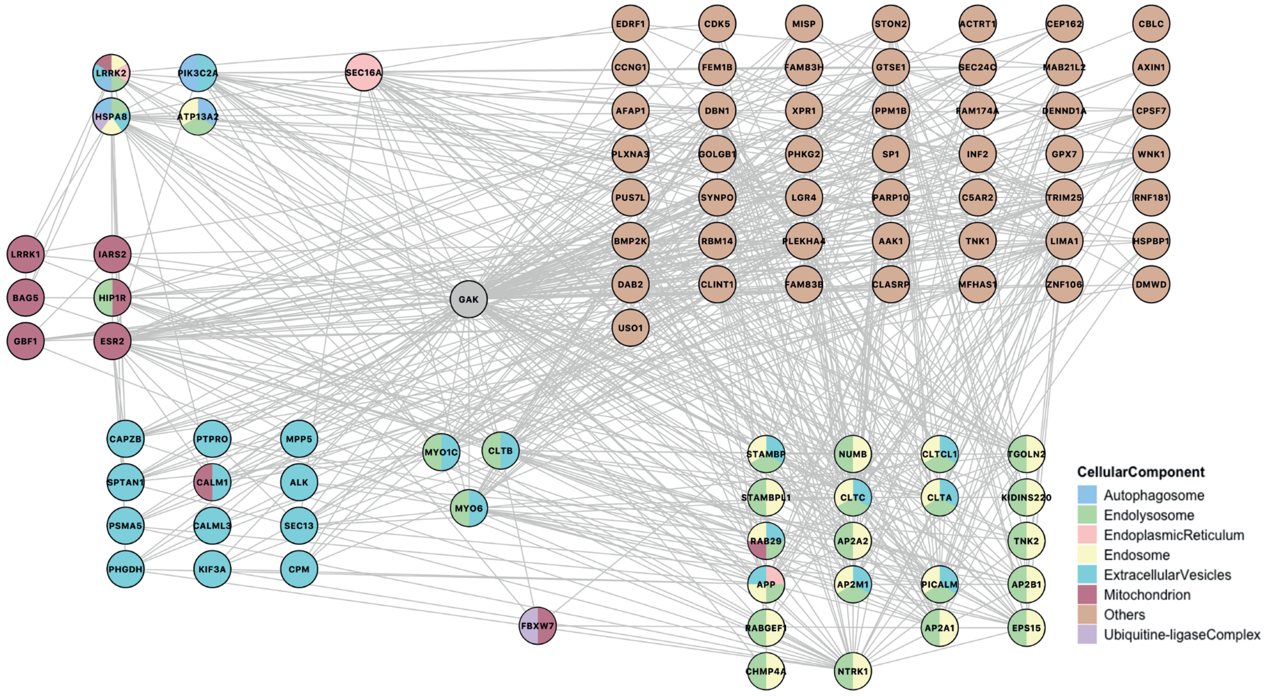


**b**

#### PRKCD Interaction Map

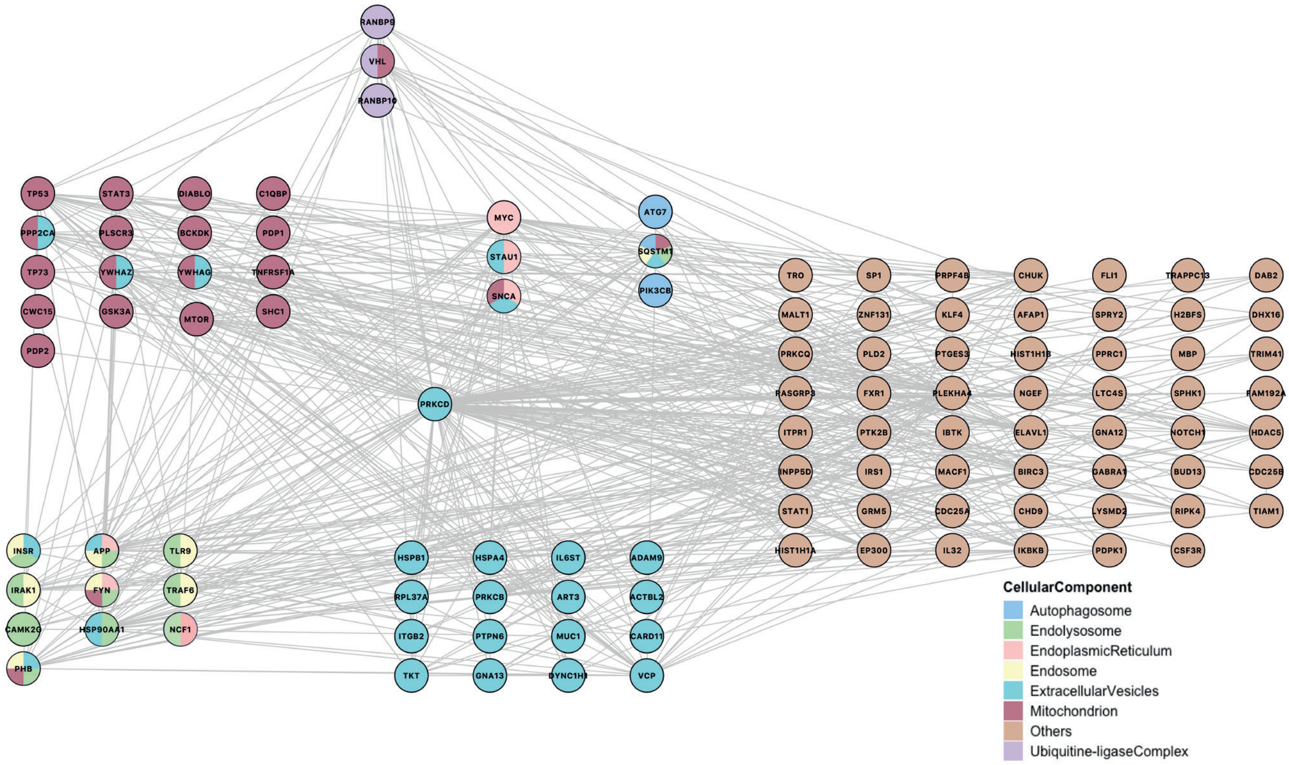


**Supplementary Fig. 2 – GAK and PRKCD Interaction Maps**

Interaction maps were generated using interaction data obtained from BioGRID (see Methods). Interactors of **a** GAK and **b** PRKCD are shown with GO analysis carried out on interacting proteins to define cellular compartment.

### Supplementary Figure 3

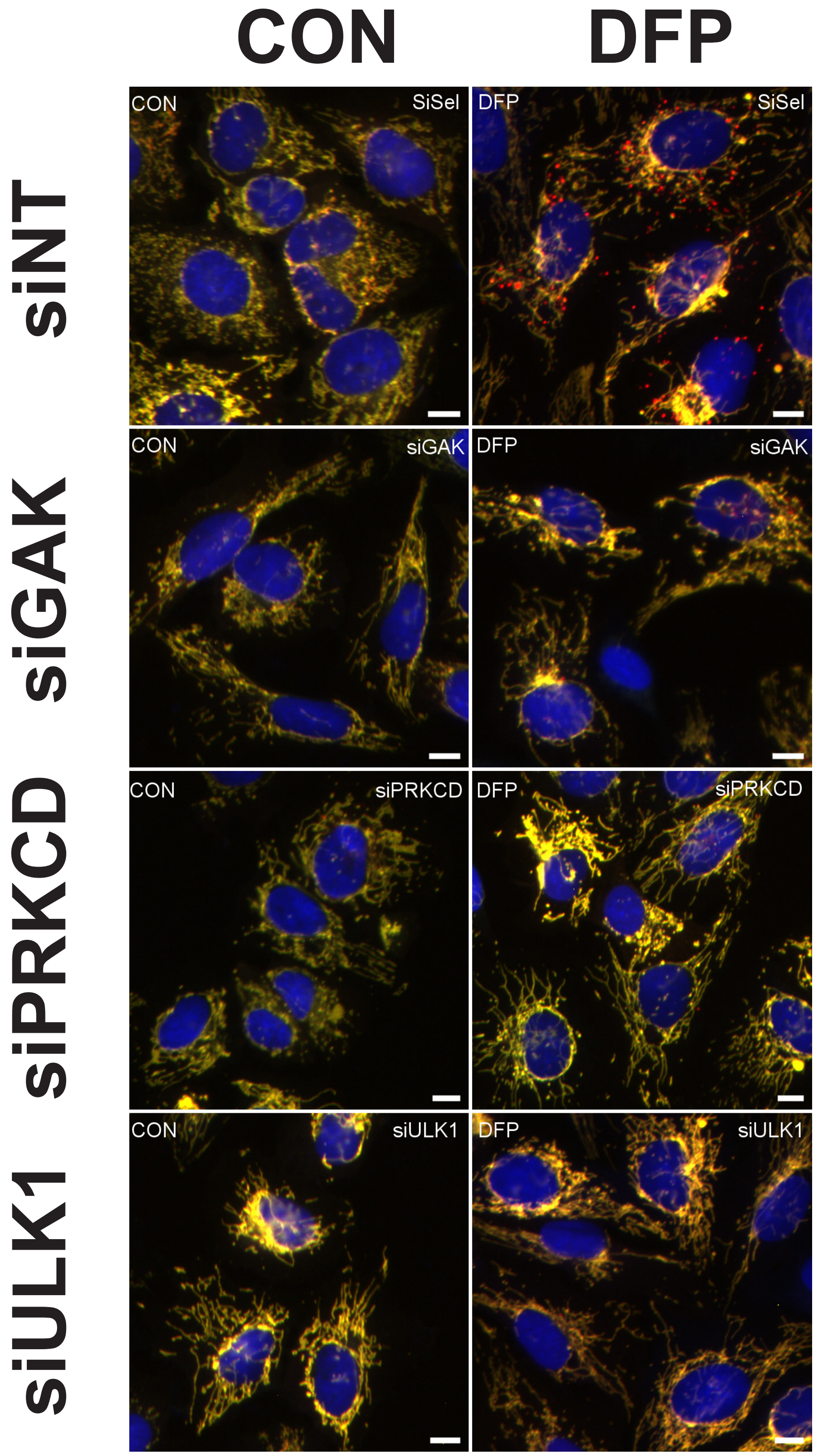


**Supplementary Fig. 3 – siRNA screen target examples**

Representative fluorescence images of siGAK, siPRKCD, siULK1 and siNT control treated U2OS cells ± 1 mM DFP for 24 h and stained with DAPI (blue). Scale bar = 10 μm

### Supplementary Figure 4

**a**

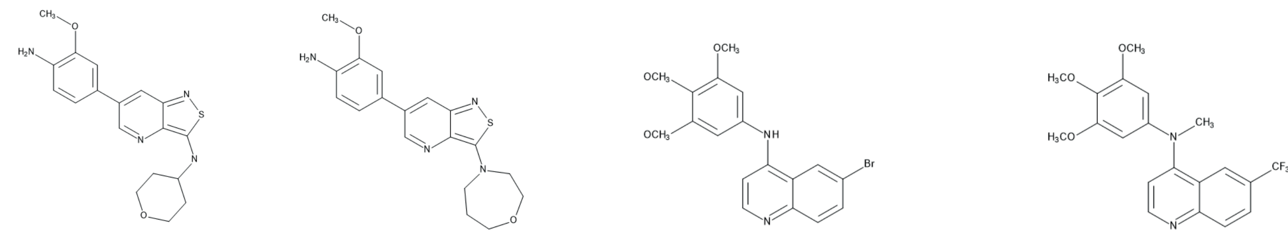

| | 12g (IVAP1966) | 12i (IVAP1967) | SGC-GAK-1 (GAKi) | SGC-GAK-1N (GAKc) |
|---|---|---|---|---|
| Kd*/Ki# | 0.08μM* | 0.19μM* | 0.0031μM* | 2.5μM# |

**b**

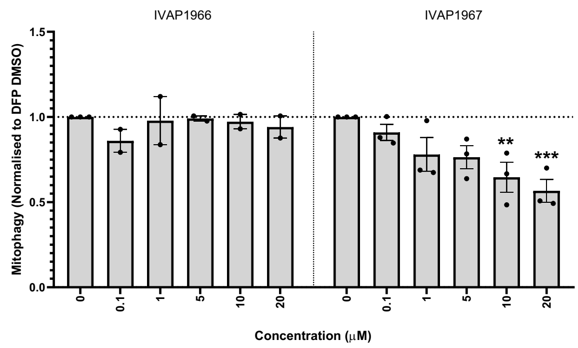


**c**

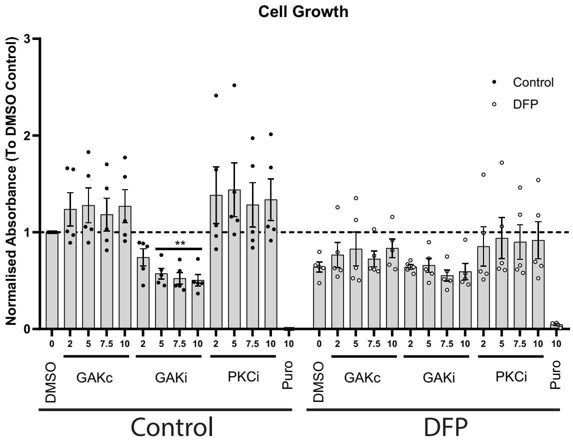


**Supplementary Fig. 4 – GAK inhibitors and effect on cellular growth**

**a** Structural comparison of different GAK inhibitors utilised in this study and their Kd/Ki values. **b** U2OS IMLS cells were treated for 24h ± 1mM DFP in the presence of indicated kinase inhibitors. Cells were fixed, imaged and quantified for red only structures and plotted relative to DFP DMSO control. Bars represent mean red structures ± SEM from n= 2 (IVAP1966) or n=3 (IVAP1967) independent experiments. Significance was determined by one-way ANOVA and Dunnett's multiple comparison test to the DMSO control. **c** U2OS cells were treated with indicated concentrations of inhibitors for 24 h prior to crystal violet staining to determine cell viability (see methods). Values represent fold change in cell number relative to DMSO control ± SEM from n=5 independent experiments. Where noted, * = p < 0.05, ** = p < 0.01, *** = p < 0.001, **** = p < 0.0001 and n.s = not significant.

### Supplementary Figure 5

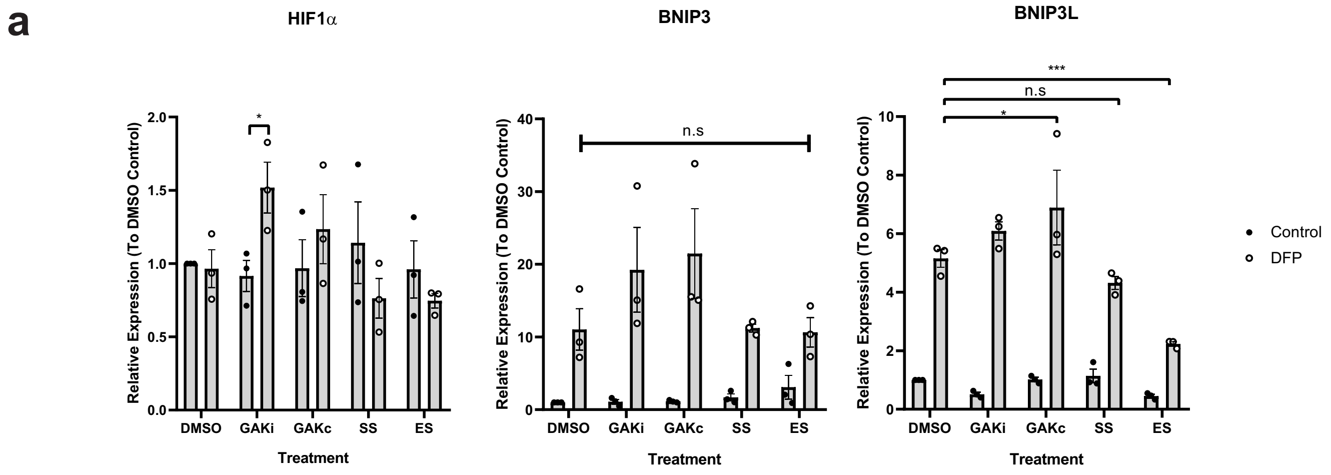


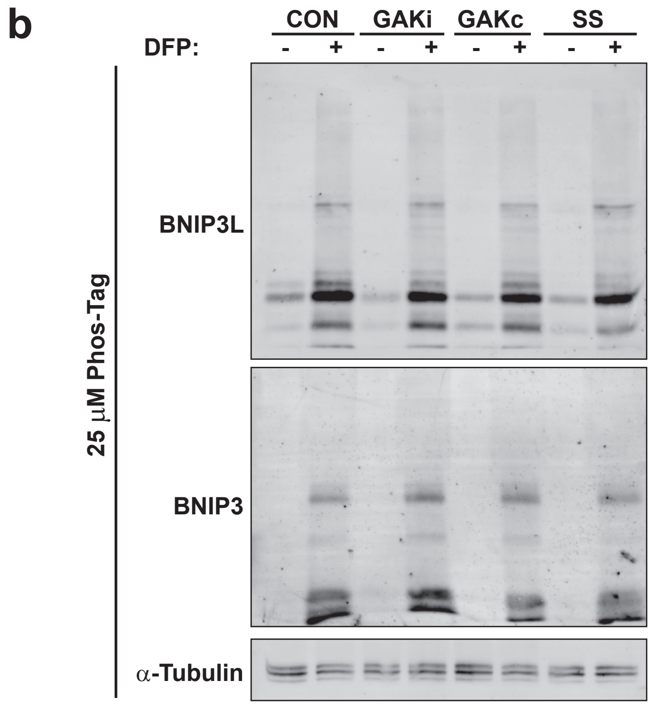


**Supplementary Figure 5 - PKCi and GAKi in HIF1α signalling**

**a** U2OS cells were treated ± 1 mM DFP for 24 h in addition to GAKi/GAKc (10 μM), sotrastaurin (SS - 2 μM), enzastaurin (ES - 2μM) or DMSO control prior to RNA isolation. qPCR analysis was carried out to determine the level of BNIP3, BNIP3L and HIF1α transcript levels after normalisation to TATA Binding box protein (TBP) followed by normalisation to the DMSO control. Values represent mean fold change in transcript relative to the DMSO control from n = 3 independent experiments ± SEM. **b** U2OS cells were treated ± 1 mM DFP for 24 h with GAKi/GAKc (10 μM each), sotrastaurin (SS - 2 μM) or DMSO control. Samples were ran on an 8 % acrylamide gel containing 25 μM Phos-Tag reagent and blotted for indicated proteins to identify phosphorylation induced band shifts[32]

### Supplementary Figure 6

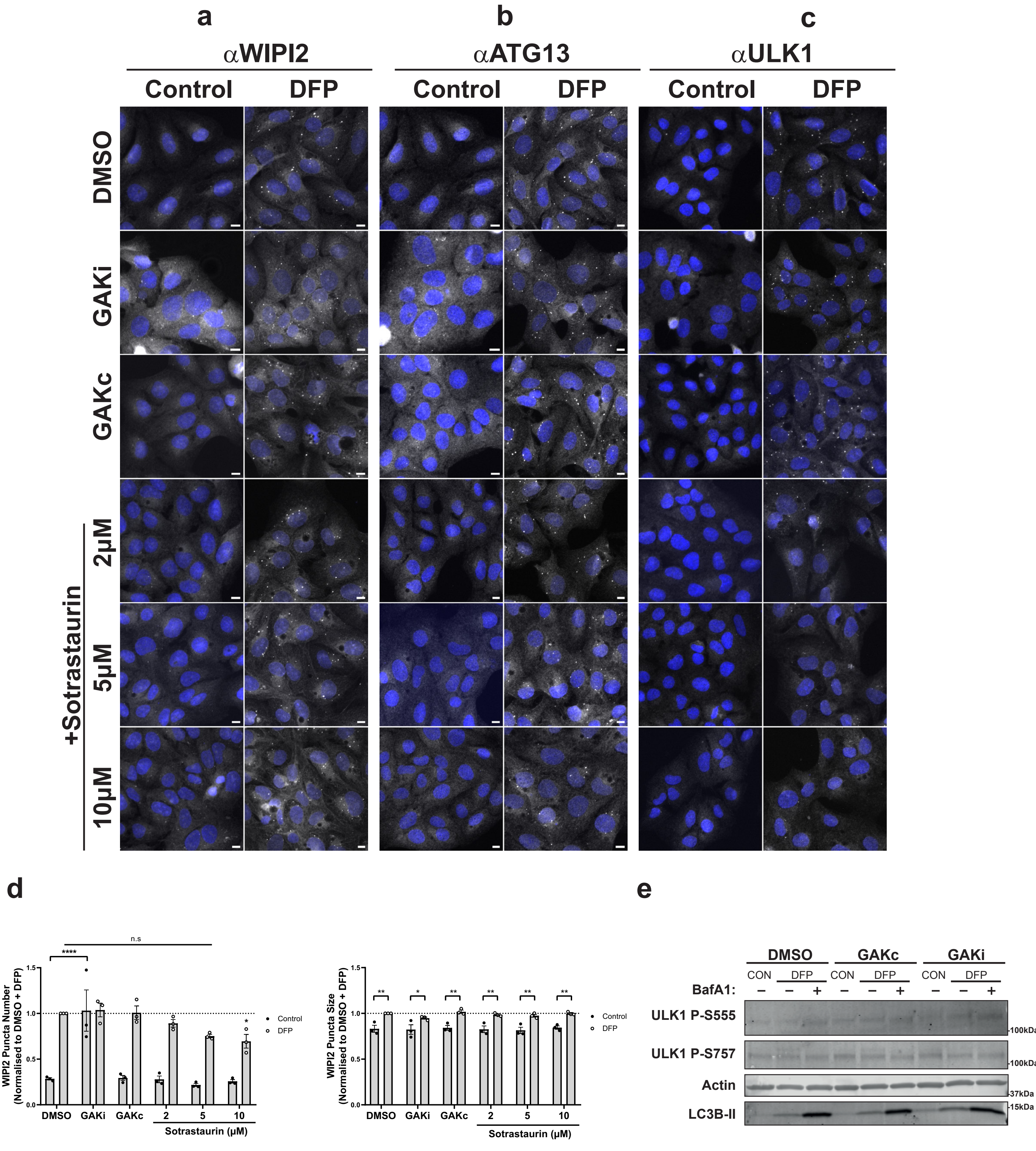


**Supplementary Figure 6 – Early autophagic machinery recruitment with GAKi and PKCi treatment**

**a-c** U2OS IMLS cells were treated ± 1mM DFP for 24h ± GAKi (10μM), GAKc (10μM) or Sotrastaurin (2 10μM) prior to PFA fixation and co-staining with antibodies directed against endogenous early autophagy markers **a** WIPI2 **b** ATG13 **c** ULK1. Representative 20x images of cells taken by Zeiss AxioObserver are shown with DAPI staining (blue), scale bar = 10μm. d Quantitation of WIPI2 puncta formed in a from n=3 independent experiments. **e** U2OS IMLS cells were treated ± 1 mM DFP for 24 h with DMSO, GAKi (10μM) or GAKc (10μM) in the presence or absence of 50 nM BafA1 for the final 16 h. Samples were western blotted for the indicated proteins.

### Supplementary Figure 7

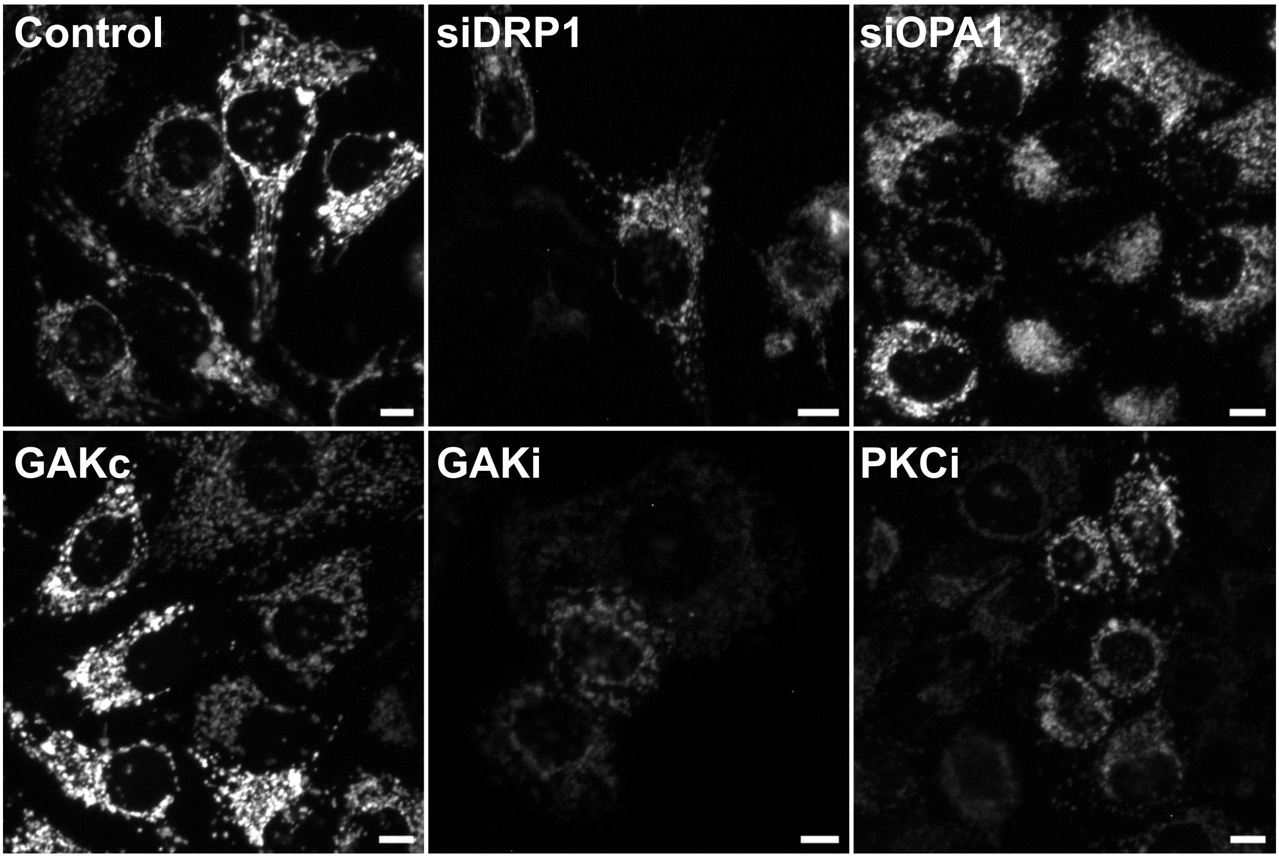


**Supplementary Figure 7 – Mitochondrial network classification**

U2OS IMLS cell mitochondrial network images (EGFP channel) used for training of machine learning classification (Fig. 7**b**). Cells were treated for 24 h with GAKi (10 µM), GAKc (10 µM) or Sotrastaurin (PKCi – 2µM) compared to 72 h knock-down of non-targeting control, siDRP1 or siOPA1. Scale bar = 10 µm

### Supplementary Figure 8

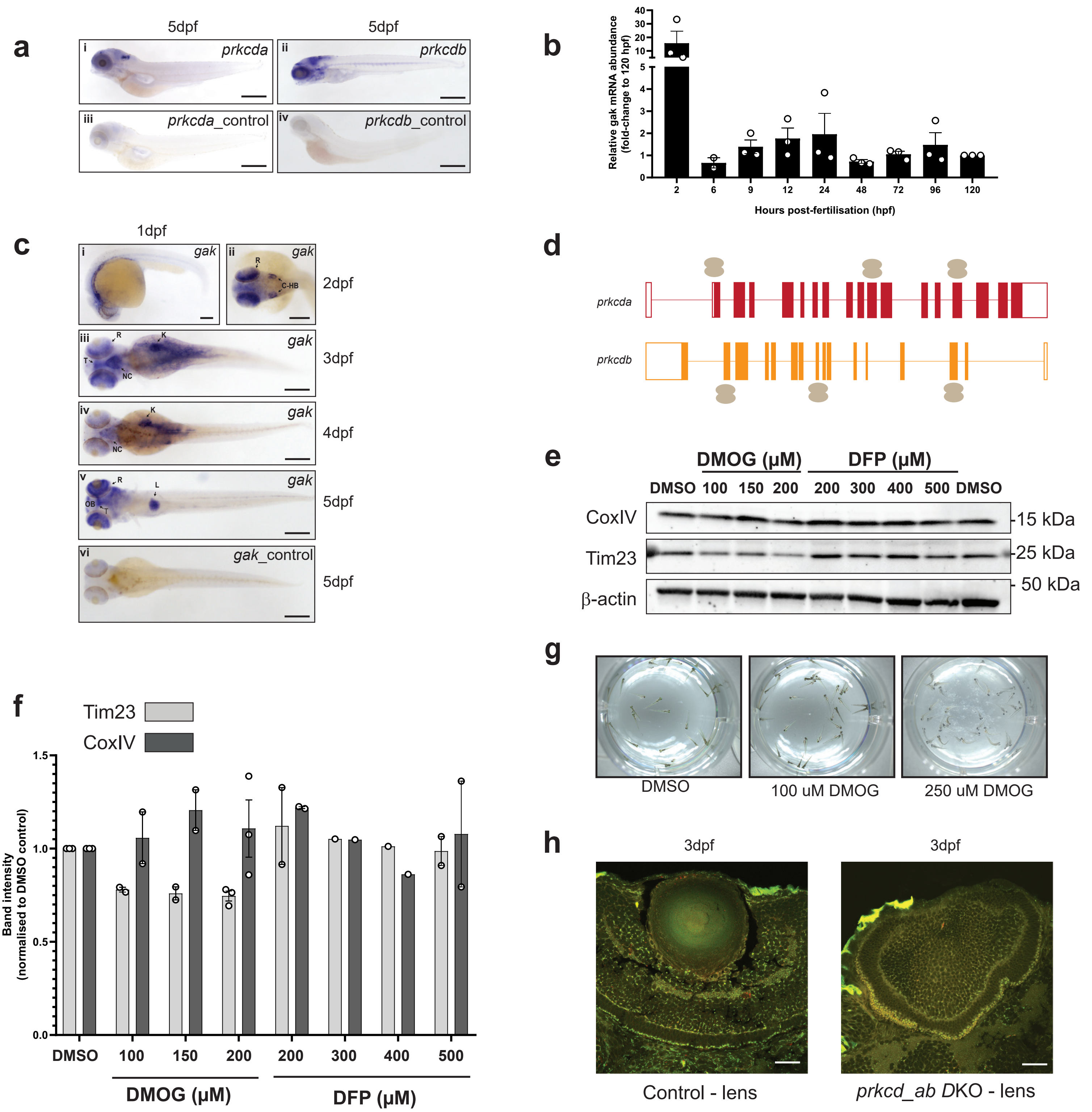


**Supplementary Figure 8 – prkcd and mitophagy in zebrafish**

**a** Spatial expression pattern of prkcda and prkcdb at 5 dpf as demonstrated by whole mount in situ hybridization (ISH) in lateral view using a 5'UTR targeting probe (i,ii) or negative sense controls (iii,iv). Scale bar = 200 μm. **b** Temporal expression pattern of gak. The graph shows the mean relative transcript abundance in whole zebrafish embryos from 2 hpf to 5 dpf from n=2 (6 hpf) or n=3 (all others) independent experiments. **c** Spatial expression pattern of gak at 1 dpf to 5 dpf as demonstrated by whole mount ISH at the indicated stages (i-v) using a 3'UTR targeting probe compared to a negative sense control probe (vi). Marked regions indicate retina (R), caudal hindbrain (C-HB), neurocranium (NC), optical tectum (T), kidney (K), olfactory bulb (OB) or liver (L). Scale bar = 200 μm. **d** Crispr-Cas9 manipulations of prkcda and prkcdb gene. Schematic description of the exon (red and orange boxes respectively) and intron (red and orange line respectively) structure of prkcda and prkcdb gene respectively. Double spherical structure with helical strands on specific boxes indicates Cas9 along with sgRNA targeting specific exons. Exon 2, 11 and 15 were targeted to create prkcda KO, whereas exon 2, 8 and 14 were targeted to create prkcdb KO. **e** Representative immunoblots of Cox IV, TIM23 and β-actin on whole embryo lysates of WT larvae treated with varying concentrations of DMOG and DFP for 24 hours at 3 dpf. Control was treated with DMSO for 24 hours. β-actin serves as the loading control. **f** Quantification of the TIM23 and Cox IV signal intensities from blots in **e** normalised to DMSO signal intensity. Bars indicate mean ± SD. **g** Representative images of control (DMSO) and DMOG treated larvae at 3 dpf. **h** representative confocal images of control (guide only) and prkcd_ab DKO transgenic tandem-tagged mitofish larvae showing the right eye. Scale bar = 20 μm.
